# Evaluating performance bias in face-to-BMI vision transformer models across diverse human populations

**DOI:** 10.64898/2026.09.02.748815

**Authors:** Jordie Hoffman, Michael Gurven, Hillard Kaplan, Jonathan Stieglitz, Benjamin C. Trumble, Bret Beheim, Paul L. Hooper, Richard B. Lee, Julia R. Phelps, Kim Hill, Brian F. Codding, Simon Brewer, Yvonne A. L. Lim, Amanda J. Lea, Ian J. Wallace, Vivek V. Venkataraman, Thomas S. Kraft

## Abstract

Computer vision models that estimate body mass index (BMI) from facial features offer a non-invasive, low-cost alternative to physical measurement, with uses in telemedicine, emergency care where a scale or measuring tools aren’t available, automated self-monitoring, and large-scale epidemiological research. Most of these models, however, are trained on government records, social media images, and celebrity photographs, sources that introduce dataset biases and fail to represent the general public. This study tests how well a face-to-BMI machine learning model generalizes across populations, specifically how morphological diversity and population-specific training data affect cross-cultural accuracy. We trained and evaluated Vision Transformer (ViT-H/14) models on paired BMI measurements and facial photographs from four Indigenous populations: the Orang Asli of Malaysia, the Ju/’hoansi of Southern Africa, the Sama residing in the Philippines, and the Tsimane of Bolivia. To evaluate how training data composition affects predictions, we compared four training strategies, from single-population models (focal models) to models trained on the full combined global dataset (global models). In-distribution training always produced the best performance.

Models exposed to a target population’s morphology, whether focal or global, consistently predicted BMI most accurately for that population. But when a target population differed from the training sample, adding more cross-cultural variation to training improved out-of-distribution predictions. Therefore, training on a population’s own data works best when that data exists, and training on data spanning a wide range of human morphology is the strongest fallback when it doesn’t. These findings suggest that while target population training data produces the most accurate results, training on datasets that capture global morphological variation substantially improves performance in unrepresented populations. Broader diversity in training data is essential for developing machine learning health tools that generalize reliably across human populations.

**Author summary:** Computer vision models that estimate body mass index (BMI) from photographs offer a promising, non-invasive, and low-cost tool for telemedicine, emergency care, and large-scale global health research.

However, the majority of published models rely on a limited number of training datasets, all drawn from urbanized, market-integrated populations with measurements that are self-reported, or estimated. While limited studies highlight risks for ethnic bias in current facial analysis models, it remains unknown whether face-to-BMI models fail to generalize across diverse global populations. To test for this, we trained a Vision Transformer model using paired photographs and BMI measurements collected by anthropologists in four morphologically and geographically distinct Indigenous populations. We found that these models are most accurate when trained on the same populations they will be assessing. When that specific data is unavailable, training on a broad cross-cultural dataset can provide a strong alternative. Our findings indicate that diverse training datasets are essential for developing accurate face-to-BMI models.

## Introduction

Computer vision models can estimate a range of anthropometric and health-related traits from facial images alone, including age, sex, and body mass index (BMI) [1–4]. Among these traits, BMI is of particular interest because it remains one of the most widely used indicators of obesity and cardiometabolic disease risk, though it is increasingly recognized as an imperfect metric when used in isolation [5–7]. Facial adiposity can be a reliable indicator of BMI [4, 8, 9] and local facial fat deposits (e.g., cheek fat) correlate closely with metabolically active visceral fat [10–12], both of which are predictors of non-communicable disease (NCD) risk. Estimating BMI from facial photographs could prove useful in settings where direct anthropometric measurements are difficult or impossible to obtain, such as telemedicine and emergency medical care [13] and large-scale population health surveillance [14]. There is also potential clinical value for patients who present infrequently for medical assessment; photographs taken throughout a patient’s lifetime could allow clinicians to reconstruct longitudinal changes in body mass. Beyond these contemporary clinical uses, applying these models to historical photographic archives also allows for retrospective analysis. This approach has precedent in economic history and human biology, where body measurements have long served as markers of health and living standards in past populations [15, 16].

While facial adiposity correlates with BMI [17, 18], population-level biological differences limit the generalizability of face-to-BMI algorithms. Populations with the same BMI often have different body fat percentages [19, 20], body fat patterning differs across ethnicities even at identical BMI levels [21–23], and baseline facial geometry varies by ancestry [24]. This morphological variance makes face-to-BMI models sensitive to domain shifts, with performance declining when models are applied to populations outside their training data [25]. Whether existing models generalize accurately across diverse populations remains unclear. Previous models have drawn on several data sets: Morph II [4, 18, 26, 27], VisualBMI [1, 14, 28–30], Bollywood dataset [29, 30], VIP Attribute [2, 18, 29–31], FIW-BMI [1, 18, 25, 26], IllinoisDOC [1, 31, 32],

MCD-rPPG [25, 33], datasets gathered from public online sources (e.g., Reddit, websites such as [www.celebheights.com](https://www.celebheights.com), [www.howtallis.org](https://www.howtallis.org) and celebsize.com) [3, 6, 25, 34], and, rarely, data gathered by the researchers themselves [17]. Relying on such narrow convenience samples creates risks of algorithmic performance bias and what Ibrahim et al. [35] call “health data poverty”: inequalities in the ability to benefit from medical innovations because the data used to develop them are not adequately representative.

Poor cross-population generalizability is a persistent problem across quantitative biology. For instance, polygenic risk scores consistently perform worse in non-European ancestry populations because the scores rely heavily on Eurocentric training data [36–39]. Facial analysis models show the same pattern: age-predicting models, for instance, have been shown to lose predictive power when applied to ancestry groups underrepresented in training data [40–42]. To determine whether face-to-BMI models suffer from this same limitation, we evaluated the cross-cultural generalization of face-to-BMI machine learning models using a dataset that spans a broad range of human morphological diversity, including disparately related populations: the Orang Asli of Malaysia (a collection of 18 ethnolinguistic groups) [43], the Tsimane of Bolivia [44], the Ju/’hoansi of Southern Africa [45], and Sama residing in the Philippines [46]. These data originate from long-term anthropological research in which paired facial photographs along with height and weight measurements were collected. Since BMI was consistently available across these diverse, non-industrialized societies, we tested whether face-to-BMI models generalize across populations. We evaluated two hypotheses regarding the impact of morphological diversity on model performance: (i) model performance for a given population will be maximized when data from that target population are included in the training set, such that non-target population training data is either irrelevant or detrimental, and (ii) model performance for out-of-sample populations will be maximized by incorporating a wider range of populations in the training set.

To test these hypotheses, we used four model configurations (Table 1): (1) focal models, trained and validated on individual populations to establish within-group predictive performance; (2) “leave-one-group-out” (LOGO) models [47] used to assess how accurately models predict an out-of-sample population when trained exclusively on other groups; (3) combination training set models, used to evaluate how incrementally adding or subtracting groups affects prediction accuracy; and (4) a global model, trained on the full diversity of available populations. Together, these four configurations directly test both hypotheses. The first hypothesis is supported if performance is best for models trained on target population data (focal & global > LOGO). The second hypothesis is supported if model accuracy increases with greater training sample diversity (global > LOGO > focal).

**Table 1.** Overview of the main model configurations. The lower panel shows the expected accuracy performance rankings under the two primary hypotheses.

| Run Type | Training Set | #<br>Runs | Purpose |
| --- | --- | --- | --- |
| Focal | Single isolated population | 4 | Evaluates performance using only local facial morphology. |
| LOGO | Three of the four available populations | 4 | Assesses generalizability to out-of-distribution groups. |
| Combination | 1 to 4 populations (all combinations) | 15 | Evaluates the incremental impact of adding groups and diversity. |
| Global | All four populations combined | 1 | Establishes baseline accuracy across multiple known groups. |
| Expected Model Accuracy Rankings For Hypotheses Tested |  |  |  |
| H1 | Focal > Global > LOGO |  |  |
| H2 | Global > LOGO > Focal |  |  |

## Materials and methods

### Dataset

#### Formation of the dataset

Measurements of height and weight, along with corresponding facial images, were compiled to form a cross-sectional dataset from anthropological field studies around the world (Table 2). BMI was calculated by dividing participant weight (kg) by the square of height (meters).

**Table 2.** Overview of study populations, locations, and data sources.

| Group | Source | Location | N | Age Range | % Male | Description | Key Cit. |
| --- | --- | --- | --- | --- | --- | --- | --- |
| Tsimane | Tsimane Health and Life History Project (THLHP) | Bolivia | 1434 | 16–81 | 49.3% | The Tsimane are an Indigenous population from the lowland Amazonian region of Bolivia in the Beni Department; their livelihood predominantly centers on small-scale shifting agriculture (horticulture), supplemented by hunting, fishing, gathering wild foods, and trade with nearby market towns. | [44] |
| Orang Asli | Orang Asli Health and Lifeways Project (OA HeLP) | Peninsular Malaysia | 1213 | 16–91 | 36.4% | The Orang Asli are the Indigenous peoples of Peninsular Malaysia. There are 19 subgroups within Orang Asli, nine of which are represented here. These groups generally practice some combination of hunting and gathering, horticulture, fishing, trade of jungle products, and engagement in wage labor. | [43] |
| Ju/’hoansi | TSpace | Southern Africa | 284 | 16–83 | 46.8% | The Ju/’hoansi are traditionally mobile hunter-gatherers who primarily rely on hunting wild game and gathering wild plant foods, with social organization based on small, fluid bands. | [45] |
| Sama | Institute of Human Origins | Southern Mindanao Island, Philippines | 88 | 16–72 | 67.0% | The Sama are traditionally characterized by seafaring nomadism, semi-nomadic, and primarily engaged in marine resource exploitation; their economic activity predominantly centers on coastal and open-ocean fishing and collecting wild foods from the local environment, supplemented by small, infrequent amounts of opportunistic wage labor. | [46] |

For the Ju/’hoansi participants, anthropometric data were originally collected by Nancy Howell and photograph data by Howell and Richard Lee, during ethnographic and demographic fieldwork among the Dobe !Kung Ju/’hoansi of western Ngamiland, Botswana, between August 1967 and May 1969 [48, 49]. Standing heights were measured on a custom-built, free-standing wooden and metal apparatus with 0.25-inch divisions designed at the University of Cape Town Medical School. Weights were recorded using an Avery butcher’s steelyard scale; subjects were weighed barefoot in minimal clothing, and the average weight of the clothing was later deducted to calculate precise weights [45].

We used data collected by the Orang Asli Health and Lifeways Project (OA HeLP) that were collected between 2020 and 2025. Standing height was measured barefoot using a stadiometer, and body weight was assessed using a digital bioelectrical impedance scale during biomedical screenings [43]. The term “Orang Asli” encompasses 19 diverse ethnolinguistic groups, nine of which are represented here. The Orang Asli are typically categorized into three major groups (Semang, Senoi, and Proto-Malay) representing general differences in language, population history, and phenotype, but they are closely genetically related and share regional geography [50–53]. We therefore evaluated them as a single, cohesive cohort to maximize available sample size, but specific ethnic groups could be examined separately in future work.

Tsimane data derive from the Tsimane Health and Life History Project (THLHP; [44]) collected between 2008 and 2011. Standing height was also measured barefoot using a portable Seca 213 stadiometer, and body weight using a Tanita bioelectrical impedance scale during medical visits [54].

Sama height and weight measurements were collected in subjects’ homes during monthly censuses of the study community in 2024 and 2025. Both height and weight were collected while subjects were barefoot, with height measured using a steel measuring tape and a flat board placed on top of the head and weight measured with a commercial digital scale after first verifying scale calibration using research assistants’ known weights that day. Photographs of subject faces were mostly collected using a smartphone camera, though three photos were taken using a Nikon digital camera.

#### Ethics Statement

Tsimane data were collected under the THLHP. Informed consent was collected at three levels: from the individual (formal written consent), at the community-level, and from the Tsimane Gran Consejo (local Tsimane governing body). Data collection was approved by the Institutional Review Boards of the University of New Mexico (#07-157) and the University of California, Santa Barbara (#3-21-0652), and the Universidad Mayor San Simon, Cochabamba, Bolivia. For Orang Asli, data were collected through OA HeLP. Informed consent was collected at multiple levels: first by describing the project to the community as a whole and seeking the permission of community leaders, and subsequently from individuals who provided formal written consent. Data collection was approved by the Medical Review and Ethics Committee of the Malaysian Ministry of Health (protocol ID: NMRR-20-2214-55565), the Malaysian Department of Orang Asli Development (permit ID: JAKOA.PP.30.052 JLD 21), and the Institutional Review Boards of Vanderbilt University (IRB #212175) and the University of Utah (protocol ID: 00197231). The Ju/’hoansi data set has been publicly archived since the 1990s in the University of Toronto’s TSpace data library, where Howell has authorized unrestricted secondary use with attribution.

Data for Tsimane, Ju/’hoansi, and Orang Asli, including identifiable photographs, were accessed for this study between August 1, 2025 and August 31, 2026. All data products generated for our analysis were subsequently deidentified and stored using ID codes.

Prospective recruitment for photographic and anthropometric data collection with Sama participants began after June 1, 2025 and continued through August 25, 2025. A bilingual research assistant read the consent form aloud, translating it into the local language, and each participant then provided written informed consent. For participants’ children aged 10 and older, the study team obtained written parental consent along with the child’s verbal assent, with a signature when the child was able to provide one.

Consent and data collection procedures were approved by Arizona State University’s Institutional Review Board (IRB #STUDY00018150).

#### Dataset Splitting

Following standard machine learning procedures, we partitioned the dataset into training (70%), validation (20%), and test (10%) subsets, drawing equally from each of the four populations. Unlike standard protocols that use the validation set for iterative hyperparameter tuning, we used ours primarily to compare performance across our different dataset configurations (i.e., focal, LOGO, combination, and global models), holding model architecture constant. The test set was reserved until the global model weights were finalized, giving us a check on whether the models generalized to new individuals rather than overfitting their training/validation distributions. Randomly partitioning the dataset presented a high risk of excluding individuals at the extremes of the BMI range from the training data. To prevent this, data partitioning was stratified by rounded BMI values (to the nearest integer) to ensure proportional representation across the BMI spectrum. For cross-validation, we divided the non-test (training and validation) data into five BMI-stratified folds, pooling the validation outputs from all folds to evaluate overall model performance [55]. Because BMI distributions vary significantly across these populations (One-Way ANOVA, F = 154.85, p<0.001; Fig 1), the model risks training primarily within each group’s mean BMI range, failing to learn how facial morphology varies at the extremes. To mitigate this issue, we implemented a weighted random sampling strategy during training. For each population, we again used rounded BMI values to the nearest integer and assigned each sample a weight inversely proportional to the frequency of its corresponding BMI bin [56, 57]. This sampling strategy increases the model’s exposure to individuals at the BMI extremes relative to their natural frequency in the data for each population.

**Fig 1.**
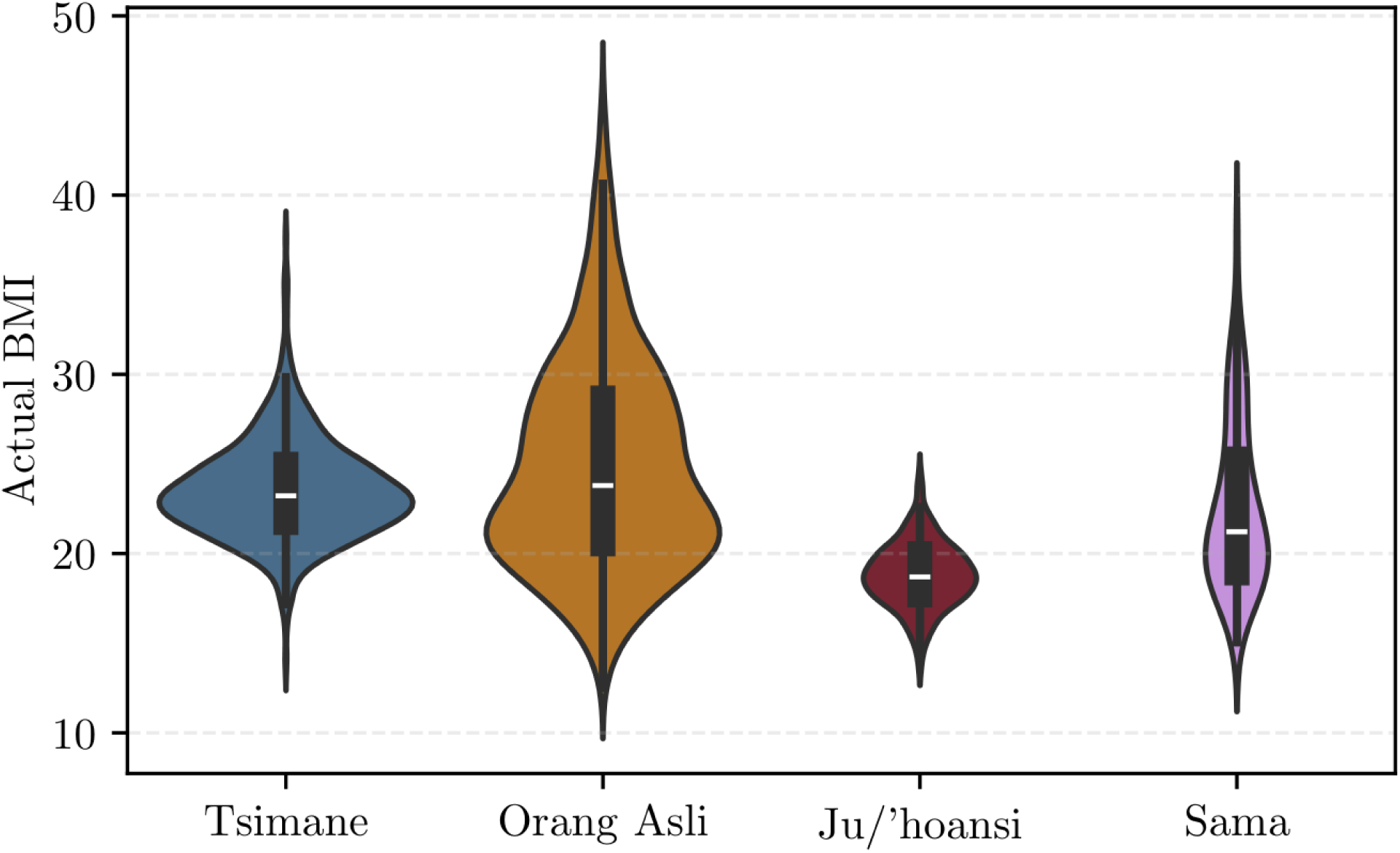
Probability density of BMI within each study population. Violin plots illustrate the probability density of BMI within each group (Tsimane n=1434, Orang Asli n=1213, Ju/’hoansi n=284, Sama n=88). Inner elements denote the median (white line) and standard deviation (SD) (thick gray bar). The Ju/’hoansi and Tsimane cohorts have the narrowest SD of 1.79 and 2.77 kg/m^2^, respectively. In contrast, the Orang Asli and Sama cohorts span wider SDs of 5.87 and 4.73 kg/m^2^.

#### Image Preprocessing

To address inconsistencies in framing, angle, and distance inherent in a dataset compiled from diverse photographic sources over different time periods, we standardized images following the principles of the ISO/IEC 19794-5 Amd 1 [58], the international benchmark used for biometric data such as electronic passports. We used the YOLO-FaceV2 model to detect faces with bounding boxes [59], and passed these to the InsightFace buffalo_l model architecture to identify a 5-point facial landmark system: left eye, right eye, nose, left mouth, and right mouth [60–69].

Following face detection, we applied a series of exclusion criteria to ensure the integrity of our dataset. Images were removed if they triggered processing errors, lacked a detectable face, or had a detected face that extended outside the image dimensions or fell within five pixels of the image’s edge. As some of these databases were not created with this type of project in mind, many records (n = 9113) were missing relevant photographs or anthropometric data. We therefore filtered the dataset based on metadata quality, discarding records with missing or duplicate information along with those involving underage subjects. We also excluded records from pregnant individuals and those with biologically implausible anthropometric measurements for the study populations, such as heights lower than 120 or greater than 200 cm, weights lower than 25 or greater than 200 kg, or BMIs lower than 10 or greater than 60. Finally, we discarded records if the BMI pre-recorded in the metadata differed by more than 5% from the BMI we calculated internally. Additional quality controls were applied, such as excluding records with a time gap of more than a year between the photograph and the physical measurements, or those with low image resolution (specifically, images where the shortest side of the original image was 180 pixels or smaller, because we found that prediction errors increased at that threshold).

To ensure horizontal eye alignment and a standardized square aspect ratio, we implemented a custom workflow that replicates Dullage’s [70] “eyelign” method. We then quantified image sharpness using Laplacian variance. Finally, we converted images to grayscale for compatibility across datasets, since some were photographed in black and white. We took this precaution because the Ju/’hoansi images were only available in black and white, and although the literature shows that models achieve virtually identical recognition accuracy across color and grayscale images, we did not want the Ju/’hoansi group to be an outlier [71, 72]. As a final quality control step, we manually inspected the photographs and their associated data, discarding 214 images where we judged the face too obscured. The final dataset consisted of 3,019 individuals (Tsimane n=1434; Orang Asli n=1213; Ju/’hoansi n=284; Sama n=88).

## Model Training

### Model Architecture

To estimate BMI from processed facial images, we used the ViT-H/14 model, a large-scale Vision Transformer (ViT) variant [73]. Unlike Convolutional Neural Networks (CNNs), ViTs partition images into fixed-size 14*×*14 pixel patches and use self-attention mechanisms to model relationships between distant regions in parallel [73]. We found that a ViT model [74] produced more accurate predictions across the range of distinct populations compared to a CNN model (Fig S1; Table S1).

We used transfer learning to adapt a model pre-trained on the ImageNet-1K dataset using the Supervised Weakly via Hashtags (SWAG) framework to the face-to-BMI task [75]. During fine-tuning, the pre-trained ViT shallow blocks were frozen to preserve foundational visual features derived from SWAG’s weakly supervised learning on large-scale image-text pairs. Following the tuning strategy demonstrated by Liu et al. [76], we selectively trained only the deeper final three transformer blocks which focus on global semantics rather than low-level textures, to encourage the model to interpret the complex facial structures unique to specific populations.

To adapt the pre-trained ViT for continuous BMI prediction, the original classification head was replaced with a task-specific multi-layer perceptron (MLP). Because overfitting is a known problem for Transformers trained with smaller datasets [77], this regression module employs a tapering architecture: it first projects 1,280-dimensional input to a 256-dimensional hidden state, and then progressively halves its width across subsequent hidden layers. This gradual reduction helps mitigate the bottleneck issues associated with abrupt dimension reduction [78]. We applied a Gaussian Error Linear Unit (GELU) activation function [79] after each linear layer and regularized the model using a dropout layer with a rate of 0.1 [80]. The model was optimized using a Mean Squared Error (MSE) loss function and the Adam optimizer with automatic mixed precision (AMP) for computational efficiency [81]. We trained the model for up to 50 epochs with an early stopping patience of 10 epochs [82].

### Evaluation Metrics

We followed previous machine learning (ML) studies that use facial images to predict BMI, which have relied on mean absolute error (MAE) and, in some cases, mean squared error (MSE) to compare model outputs [1, 2, 4, 6, 14, 17, 18, 26–32]. MAE measures the average magnitude (absolute value) of errors between observed and predicted values. However, if a group has low BMI variability, a model can achieve a low MAE simply by predicting the group’s average BMI for every individual. In this scenario, the model appears accurate, but it has not actually learned how facial features change as body weight increases or decreases [83]. To avoid this limitation, we supplemented our analysis with additional statistical metrics.

To maintain cross-group comparability despite differences in the population’s BMI variance, we calculated Relative Error (RE). This metric normalizes the Root Mean Squared Error (RMSE) by the sample standard deviation of the true BMI values, denoted by *s_y_*:

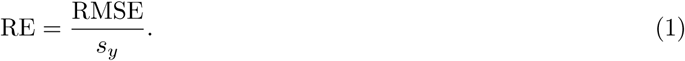

To further assess model performance, we derived the coefficient of determination (*R*^2^) and the regression slope (*β*) from models fitting observed versus predicted values. Alongside linear and residual plots, we used these metrics to evaluate how accurately the model’s predictions captured the true variation in BMI across the dataset populations.

Because ML models cannot reliably estimate the BMI of an individual outside of the training dataset’s range (Fig S2), we restricted prediction analyses to only consider individuals with a measured BMI within the training range. For the combination set analysis, this restriction allowed for a clear comparison between the performance improvement gained by including target population data versus the improvement gained by adding non-target data from the same BMI range. Consequently, for the Orang Asli (which had the lowest and highest BMIs), outlying data points below 14 and above 35 BMI were excluded. This ensured the analysis solely evaluated BMI ranges containing individuals from at least two different demographic groups.

We evaluated the model’s clinical utility by assessing residual errors against established clinical thresholds.

Following clinical guidelines that define changes in body weight of 5% and 10% as clinically meaningful thresholds [7, 84, 85], we measured the proportion of model prediction residuals falling within both these boundaries. To determine whether model errors were large enough to clinically miscategorize individuals, we also compared observed and predicted BMIs across standard World Health Organization (WHO) classifications (i.e. Underweight, Normal, Overweight, Obese I–III) [86] (Table S2).

### Generalizability and Model Comparison

Generalizability of the global model was tracked by categorizing validation groups as in-distribution, those included in the training phase, or out-of-distribution (OOD), those held out until the validation phase. For direct performance comparisons between the LOGO and focal models, the concordance correlation coefficient (CCC) was utilized to measure the agreement between paired residual sets *x* and *y*:

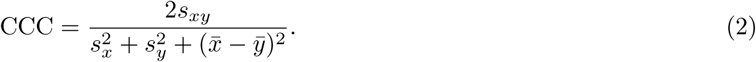

Additionally, a win rate percentage was calculated to determine the frequency with which the LOGO model achieved a lower absolute error than the focal model. This metric represents the percentage of individual observations where the absolute prediction error of the LOGO configuration was less than that of the focal model.

## Results

### Focal versus LOGO Models

We predicted that if accurate BMI estimation relies on a population’s specific facial morphology, focal models would consistently outperform LOGO models. Consistent with this prediction, omitting the target population from training increased absolute and relative estimation errors for the Orang Asli, Ju/’hoansi, and Tsimane (Fig 2 a-b; Table 3). Focal training within the target population therefore consistently outperforms out-of-sample prediction for these three populations. For the Orang Asli and Tsimane cohorts, the LOGO configuration had lower explained variance. The LOGO model slope was closer to zero for the Orang Asli cohort (*β* = 0.28) than the focal model (*β* = 0.65), while the Tsimane slope remained practically unchanged (LOGO *β* = 0.40; focal *β* = 0.40). Focal models also achieved higher win rates (by an average margin of 14.3%) and stronger concordance (CCC = 0.53–0.67) for the Orang Asli, Ju/’hoansi, and Tsimane cohorts (Fig S3).

**Fig 2.**
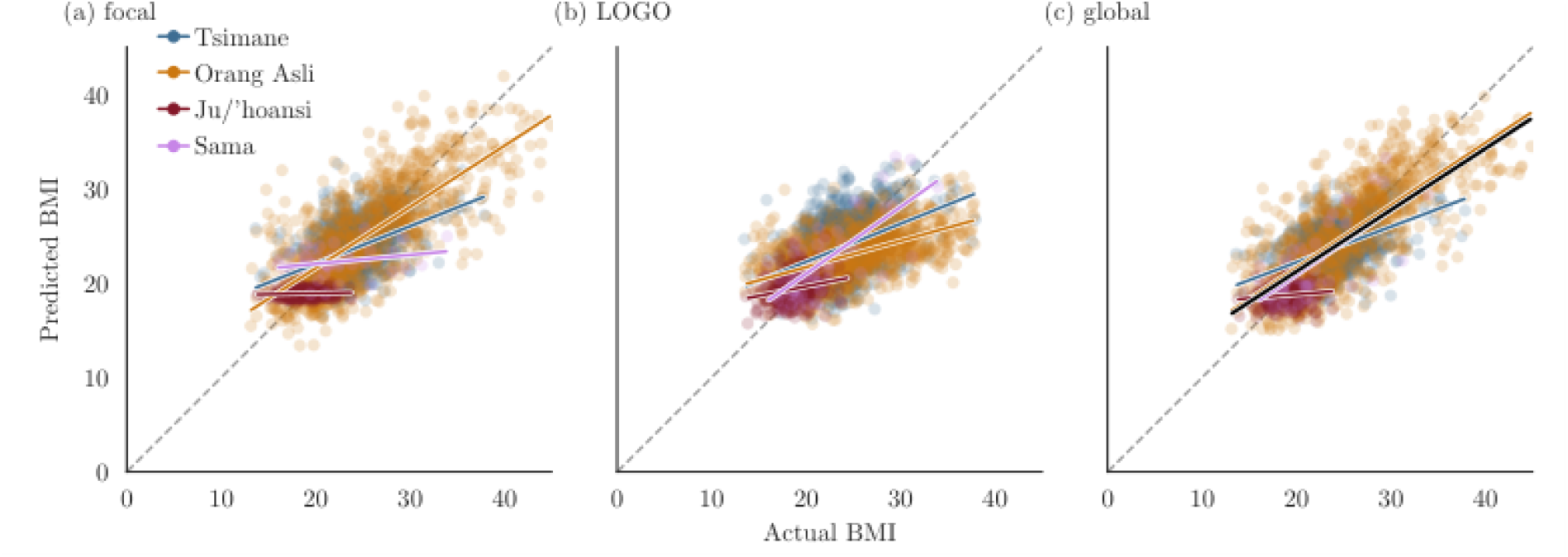
Predicted versus actual BMI across model configurations. Predicted versus actual BMI across (a) focal, (b) LOGO, and (c) global models. Solid lines indicate the linear regression fit for each demographic group; the dashed diagonal is the 1:1 perfect prediction line.

**Table 3.**
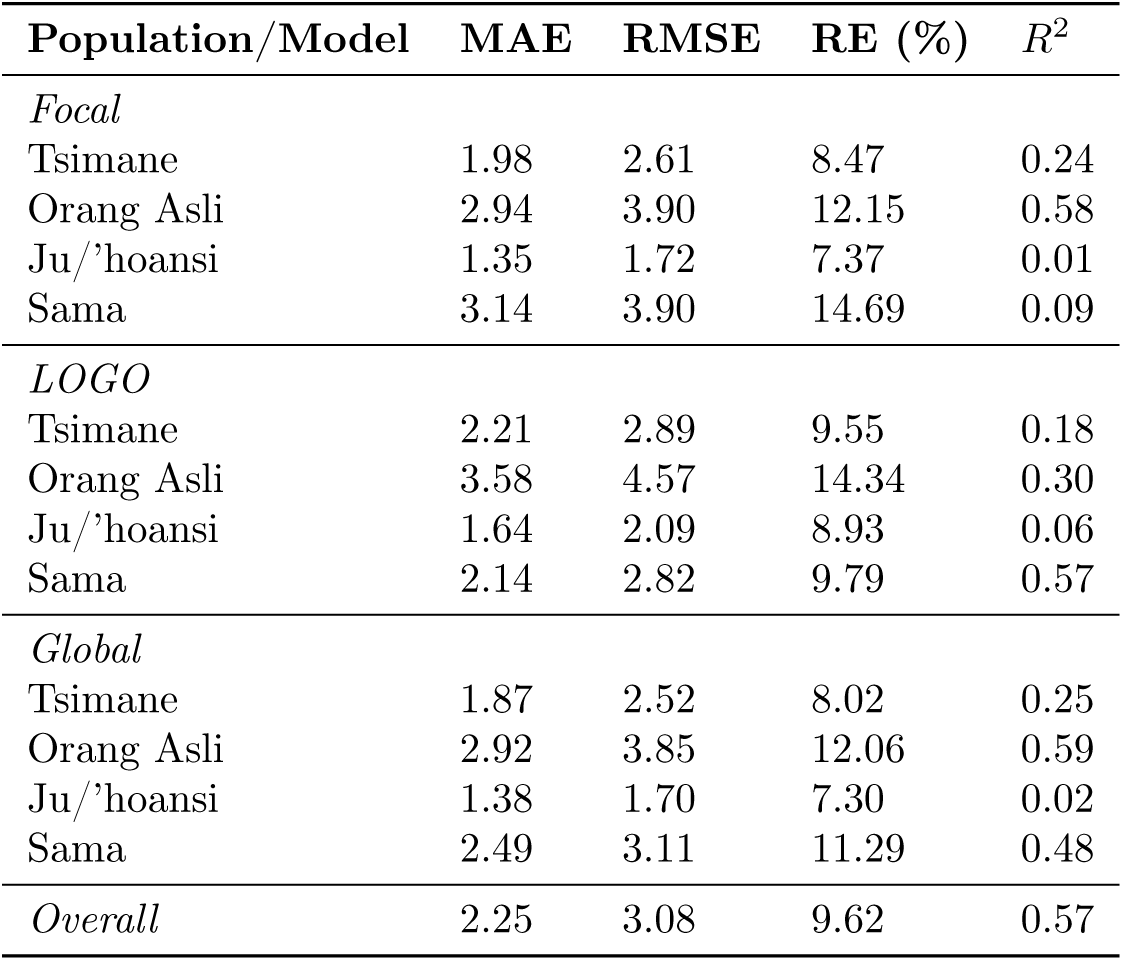
Summary of model prediction metrics by run type for each of our four populations.

There were several minor exceptions. For example, the Sama LOGO model yielded lower absolute and relative estimation errors, higher explained variance, stronger concordance, and a steeper positive slope than its focal counterpart. For the Ju/’hoansi cohort, the LOGO configuration also yielded higher explained variance and a steeper slope than the focal model, despite generating higher estimation errors. Additionally, LOGO models generated lower estimation errors than focal models from the 70th–99th percentile of the validation population’s BMI range for every group except the Orang Asli.

### Combination Training Set Models

To quantify how incrementally adding or subtracting groups affected prediction accuracy, we trained models on every combination of our four groups. For in-distribution predictions, increasing training diversity decreased absolute errors and increased explained variance (Figs 3 and 4). OOD predictions exhibited similar trends, though with greater absolute error than in-distribution predictions at each level of diversity (Fig 4). This implies that when in-distribution training is not possible, a model still benefits from training on more diverse populations.

**Fig 3.**
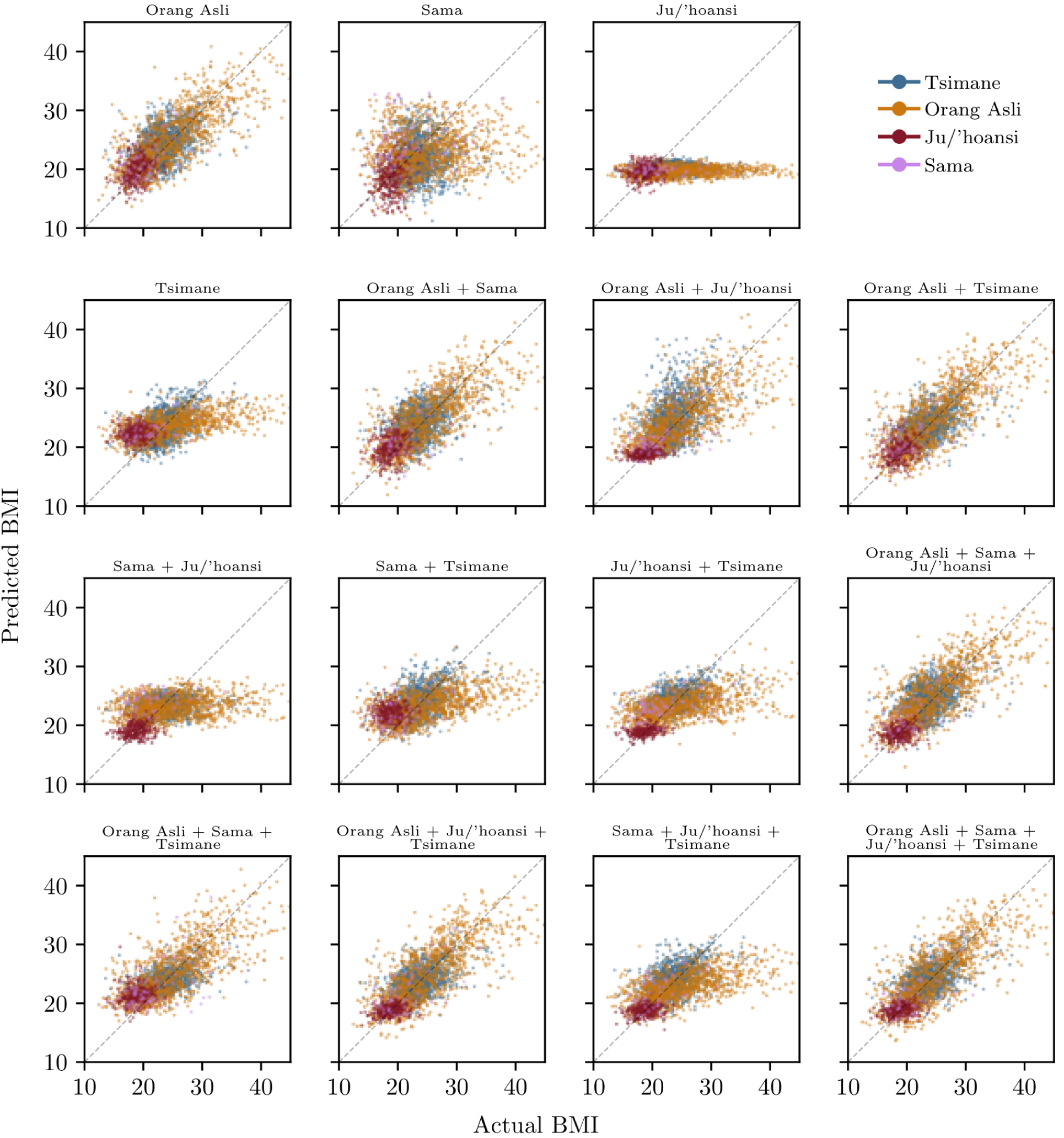
Predicted versus measured BMI for every population combination. Scatter plot comparing predicted versus measured BMI for every population combination used to train the model. Each of the fifteen panels is a scatter plot of predicted BMI against measured BMI, for one training-set composition; the panel title states which of the four populations were included in the training data. Within each panel, every individual in that model’s evaluation set is plotted and coloured by population using the colour key shown in the top-right panel, so a given colour always denotes the same group. The dashed diagonal is the line of perfect prediction.

**Fig 4.**
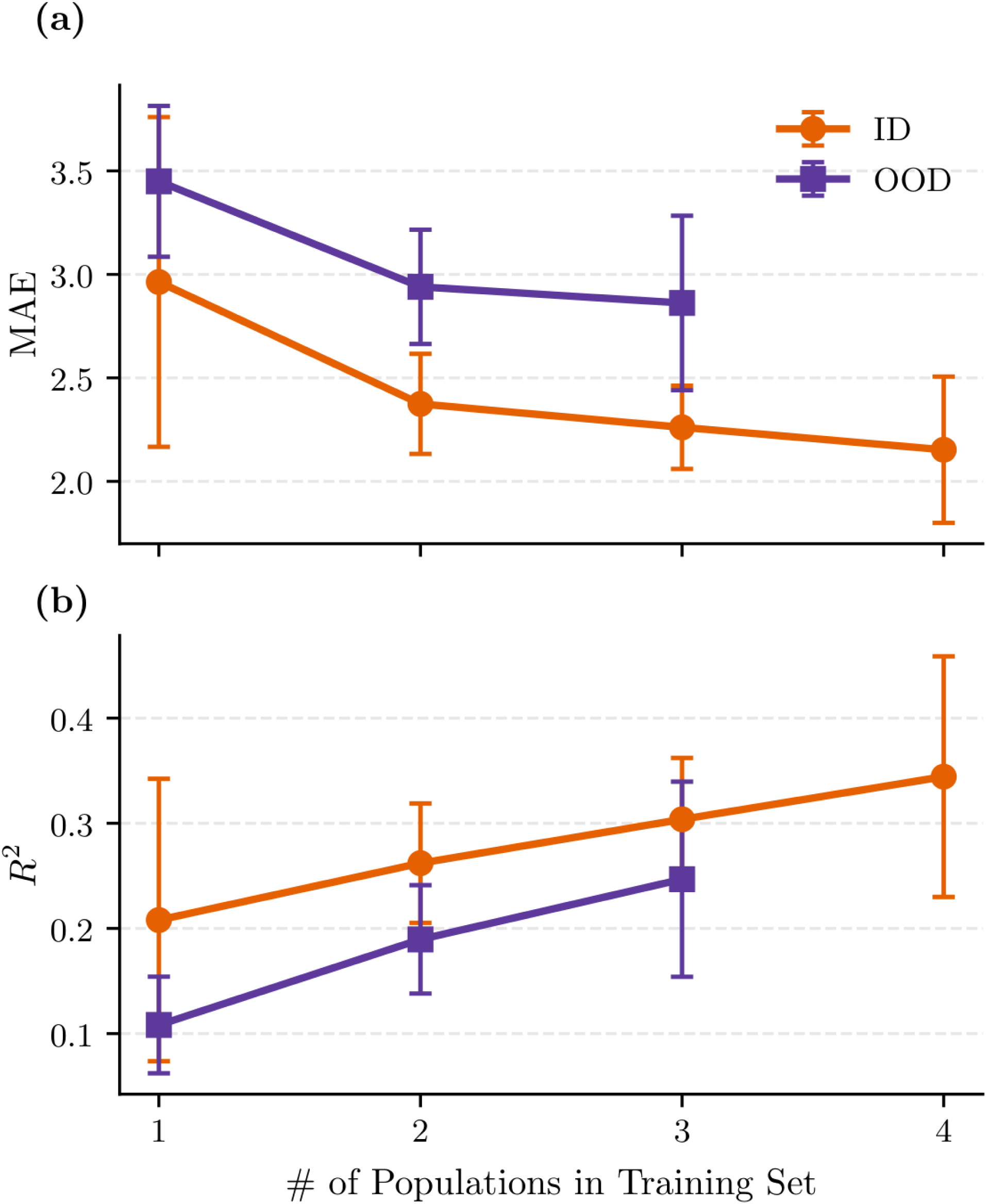
Model performance trajectories as a function of training set diversity. The panels display MAE and *R*^2^ evaluated against the number of populations included in the training set. Data points represent the aggregated mean performance across all possible combinations of the four populations. Error bars represent *±*1 SE across all population–model combinations at each training-set size. Performance is stratified by prediction status: ID evaluations are plotted as orange circles, and OOD evaluations are plotted as purple squares. Increasing training-set diversity generally decreases absolute prediction errors and increases explained variance for both ID and OOD predictions, with the largest gains occurring at lower levels of diversity.

### Global Model

In the global model, a population’s BMI variance drove both prediction error (MAE) and model fit (*R*^2^) in the same direction (Fig 2; Table 3). The global model made the largest errors on the Orang Asli and Sama cohorts, which had the most variable observed BMI, yet these groups yielded the steepest slopes and highest *R*^2^. Conversely, the model made the fewest errors on the Ju/’hoansi and Tsimane cohorts, which had the least variable observed BMI, but these groups produced the lowest slopes and *R*^2^. The deviation of an individual’s observed BMI from the median of their group positively predicted the absolute prediction error of the model (*β* = 0.32, p < 0.001). However, when evaluating all populations combined, the global model’s overall regression slope and *R*^2^ exceeded the individual metrics for the low-variance Tsimane (by 0.27 in slope and 0.32 in *R*^2^) and Ju/’hoansi (by 0.57 in slope and 0.55 in *R*^2^). Fig S4 further explores how population variance affects model generalizability.

When predicted and actual BMIs were mapped to WHO categories, the model demonstrated higher categorical agreement for individuals from populations with a low variance in BMI (Table S3, Fig 5). As a person’s BMI diverged from this population median, absolute errors increased significantly (overall *β*=0.32, p < 0.001, *R*^2^ = 0.21), decreasing categorical accuracy at the extreme ends of the spectrum. However, a tight BMI distribution did not always guarantee categorical success. For example, despite the Ju/’hoansi having the highest continuous accuracy of any group (42.9% of predictions falling within a 5% relative error), their categorical accuracy was disproportionately lower (57.9%) than most other populations, with worse WHO categorical accuracy. This discrepancy likely occurs because their mean BMI (18.8 kg/m^2^) sits near the underweight category threshold (18.5 kg/m^2^), meaning even highly accurate predictions with minor numerical errors frequently cross the boundary and result in categorical misclassifications.

**Fig 5.**
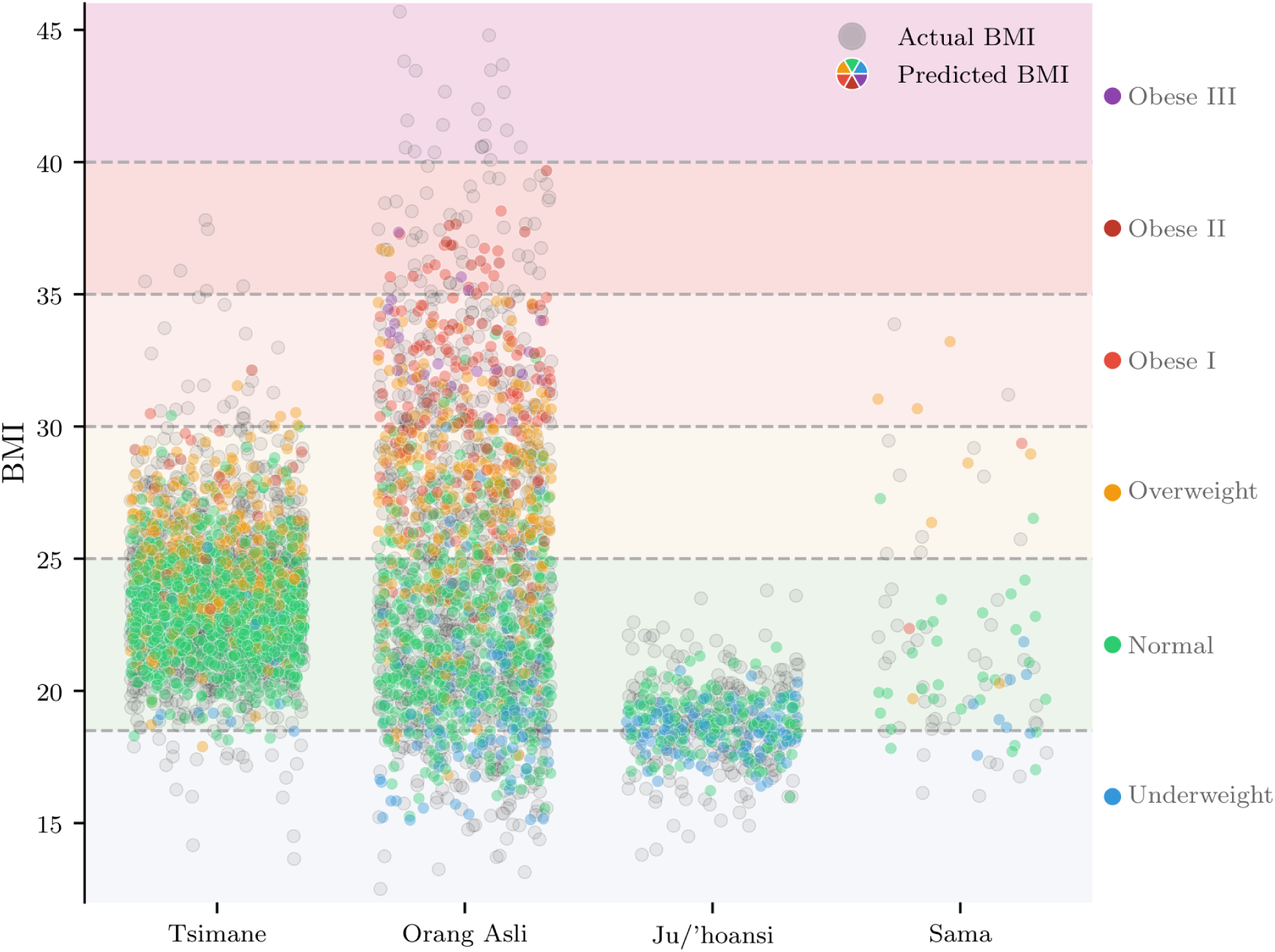
Distribution of predicted BMI across demographic groups. Overlaying predicted WHO category backgrounds. Points are colored according to the subject’s true WHO classification.

We evaluated whether predictive error was affected by age, sex, population, facial geometry, and image quality. Overall, the global model yielded lower predictive errors for males than for females (a difference of -0.84 MAE). Greater observed deviation from the median BMI was significantly associated with increased prediction error (slope = 0.09, p < 0.001; Fig S5). Differences in age, facial structure (e.g., face width-to-height ratio and face angle), image brightness, and interaction between age and sex did not affect the model’s accuracy (Fig S6). Conversely, high face detection confidence was associated with the lowest prediction errors (*≤*5%).

When we evaluated the global model weights on the withheld test subset, the MAE and *R*^2^ were generally the same as the global model for the Tsimane and Orang Asli, but were improved in the test set for the Ju/’hoansi and Sama (Fig 6). The test set produced error variance which peaked at the 25–30 observed BMI range (Fig S7). The validation set produced wider error variance at the extremes of the BMI distribution, stemming primarily from the Orang Asli and Ju/’hoansi cohorts.

**Fig 6.**
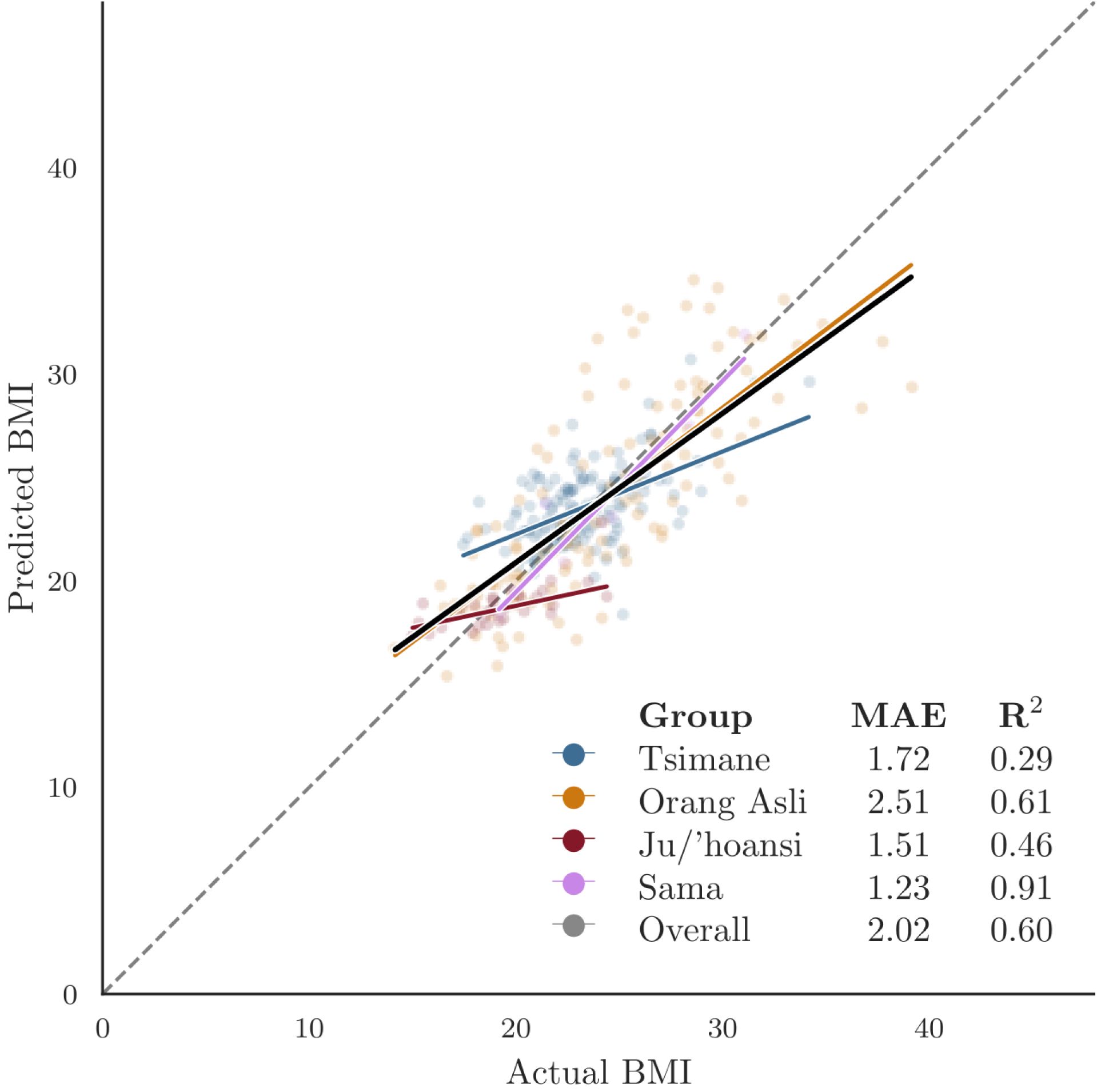
Predicted versus actual BMI on the held-out test set. Predicted versus actual BMI across our four population groups using our 10% hold-out test set. Solid trendlines show the mean predictions of the five cross-validation folds and overall prediction trendline.

### Comparison of model performance

To evaluate relative performance across our four populations, we compared the absolute errors of predictions from the focal, LOGO, and global models (Table 3). For the Orang Asli, Ju/’hoansi, and Tsimane, focal models yielded nearly identical absolute errors to global models. Conversely, omitting the target population from the training dataset (i.e., LOGO) increased the absolute error for these groups compared to their global and focal counterparts (Fig 7). Consistent with our expected model outcomes (Table 1), these findings imply that predictive accuracy is highest when models are trained in-distribution. The Sama cohort was the sole exception to this trend, as their LOGO model yielded lower estimation errors than their focal model.

**Fig 7.**
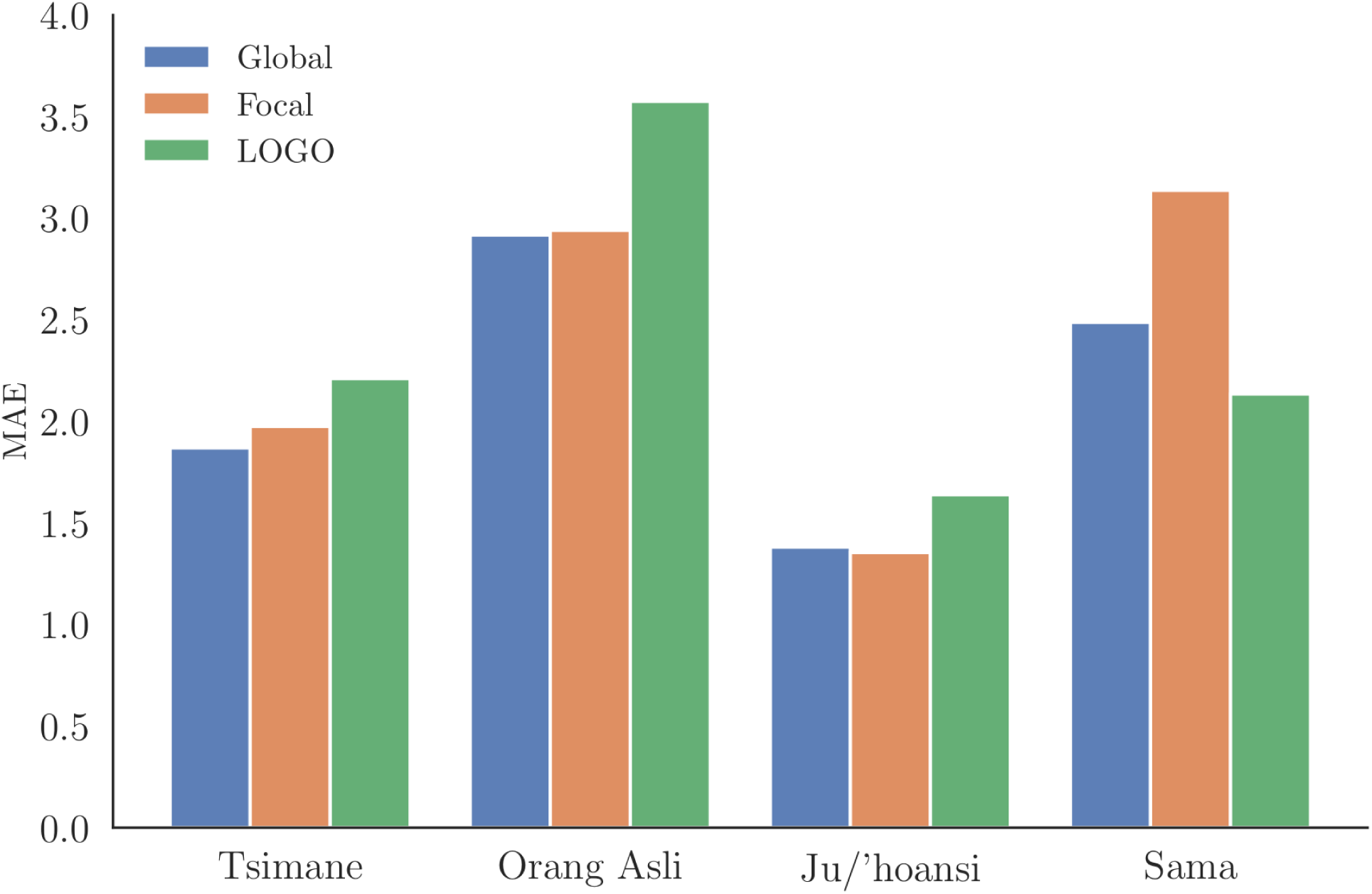
Predictive accuracy (MAE) of global, focal, and LOGO models across populations. Grouped bar chart comparing the predictive accuracy, measured by MAE, of our three models (global, focal, and LOGO) across our four populations. Each population category features a cluster of vertical bars, with each bar representing a specific model’s MAE score for that population, allowing for direct comparison of different training results within each group.

## Discussion

### Training on Included Groups Maximizes Accuracy, While Diversity Improves Predictions on Held-Out Groups

Predictive accuracy in our face-to-BMI models depended on two things: whether population-specific training data were available, and whether the training set captured universal physical cues from broad morphological diversity. Hypothesis 1 predicted that models trained on the target population, either alone (focal) or as part of a larger set (global), would outperform models trained without it (LOGO). Our results broadly supported this: global and focal models were functionally equivalent across most populations in terms of MAE, while LOGO model performance was generally poorer (Table 1). Hypothesis 2 predicted a stricter ordering, with global models outperforming LOGO models, which would in turn outperform focal models, since broader diversity should extract more generalizable physical cues. Our results only partly supported this. The most diverse (global) model’s predictions matched, rather than exceeded, the accuracy of population-specific focal training. This equivalence is notable because such models typically shrink toward the mean, so similar focal model accuracy shows that the global model training retains representation of minority populations. LOGO models outperformed focal models only in specific cases (the Sama cohort, and the upper BMI range for most groups; Fig S3). Broad diversity was therefore beneficial, but models still performed best when evaluated on populations explicitly represented in their training data (i.e., for in-distribution populations). While training exclusively on a target population (i.e., focal models) minimized absolute prediction errors for groups like the Ju/’hoansi (Fig 7), our global model had marginally better MAE, explained more variance, and prediction slopes closer to 1 across most groups.

Omitting a target population generally degraded model performance, lowering the model’s ability to keep predictions within the 5–10% error boundaries required to be clinically meaningful (Table S4). The Sama were the exception: their focal training sample was too small for the ID focal model, and its inclusion may therefore have hindered prediction from the global model relative to generalizing from the other diverse groups alone. Consequently, the most reliable path to accurate face-to-BMI prediction in populations not currently represented in large databases is to include target population images directly in model training. Most existing large databases, however, are convenience samples built from celebrity photographs, administrative records, or self-reported online data rather than direct population sampling, so models trained on them are unlikely to generalize well to populations these sources do not capture. Without local representation or highly diverse training sets, facial-analysis models lose prediction accuracy and risk encoding inequities by race, ethnicity, or sex into medical care [27, 87–89].

### Effects of Image Quality, Age, Sex, and Ethnicity on Predictive Error

A smaller, multi-population algorithm maximizing training on morphological diversity achieves a lower prediction error (ΔMAE = 0.40) than a large, single-population framework maximizing training on BMI density for a withheld target group. This suggests that broader human diversity in training improves generalizability more than sample size alone (Fig S8). Superior performance for in-distribution populations is expected given population-level differences in baseline facial geometry [24, 40] and fat patterning [21–23]. Our results also show, however, that incorporating more diversity into the training data still achieves reasonable performance (e.g., MAE < 0.13) when in-distribution training is not possible. Larger training datasets spanning a broader range of human diversity may therefore make generalizable, clinically meaningful (error within 5–10% boundaries) predictions feasible for OOD populations.

BMI cannot separate muscle from fat, and the ratio between the two shifts with sex, age, and genetics [20, 90, 91]. This distinction is relevant in populations like the Tsimane, where some men maintain a very lean appearance but have a high BMI due to dense muscle mass. Consequently, our current models cannot resolve whether predictive accuracy relies on how facial fat accumulates as BMI rises, or on how baseline facial structure varies across populations independent of BMI. Isolating these two signals will require alternative measures of adiposity [5]. Bioelectrical impedance estimates of percentage body fat exist for the Orang Asli and Tsimane cohorts in our study [43, 44], though testing our predictions against these estimates falls outside the scope of the current methodological paper. Future work should compare facial-morphology predictions to these bioimpedance estimates, which would help disentangle these signals and clarify whether the model is tracking fat deposition, population-specific facial geometry, or their interaction.

In our global model, absolute prediction error was driven by individual-level factors rather than image quality. Predictive error increased as an individual’s observed BMI diverged further from the sample median (*β* = 0.32, p < 0.001; Fig S5). Other face-to-BMI algorithms show the same pattern, largely because available training datasets underrepresent underweight and obese individuals [17, 18, 26]. Consistent with Siddiqui et al. [27], the global model also produced higher predictive errors for females than for males (Fig S5); this sex-based disparity is statistically significant given the large sample size, but amounts to an average difference of only 0.84 BMI between males and females, well below the 5% body weight fluctuation threshold that medical guidelines treat as clinically meaningful [7, 84, 85]. Photograph quality metrics did not meaningfully influence predictions (Fig S6).

### Broader Implications for Clinical and Demographic Applications

While medical practitioners and diagnostic tools frequently rely on race or ethnicity as an accessible proxy for clinical decision-making [92–94], doing so can obscure true physiological drivers of disease and lead to misdiagnoses [95–97]. Our models never received a race, ethnicity, or population label of any kind, yet still performed best on whichever population they were trained on. This suggests these models are picking up directly on facial morphology, and that it is a more precise signal than race or ethnicity. For clinical deployment, a model’s training data – whether drawn directly from the target population or from a sufficiently diverse range of populations – must adequately represent the population it will be used on.

### Limitations

Throughout all model iterations, model performance was poor for the Ju/’hoansi. While population-level MAE was consistently low, the predictions yielded relatively flat regression slopes, suggesting that the model failed to capture within-population differences. Our supplementary analyses indicate that low slopes are best explained by the Ju/’hoansi’s narrow BMI distribution, which is likely due to their lack of market integration when the measurements were taken (Fig S9). When we restricted other populations to that same narrow BMI window, they showed the same flat-slope pattern under LOGO models, which points to training density rather than Ju/’hoansi phenotype or photograph quality as the driver (Fig S9; full analysis in Supporting Information). Some of this may also reflect real non-linearity in how fat accumulates across the BMI range, a possibility we cannot yet separate from the training-density explanation. The Ju/’hoansi photographs are also unique in being scanned from 1970s film rather than digital images taken after 2000; because ViTs rely less on local texture than CNNs do, this is unlikely to be the main cause of the Ju/’hoansi’s poor performance [98], though how analog and digital image sources interact with ViT training remains understudied. Field conditions introduce other noise: inconsistent backdrops, angles, lighting, and non-facial features like head coverings, which we partly addressed with eye alignment but could not fully mitigate.

An original goal of this research was to examine whether BMI models trained solely on existing datasets, focused on populations from large-scale, market-integrated societies, can generalize to Indigenous populations not represented in those datasets. However, the available datasets used to train previous models rely on unverified, self-reported measurements extracted from online forums or celebrity-tracking websites. These sources introduce risks of reporting errors and temporal mismatches between a photograph’s capture date and the subject’s recorded weight. Even formal datasets compiled from law enforcement arrest records, such as the widely used MORPH-II database, carry risks of self-reporting and temporal inaccuracies. As Bingham et al. [99] note, most demographic data gathered for these mugshots were self-reported at the time of an arrest. Their audit of the MORPH-II dataset revealed extensive longitudinal inconsistencies across multiple entries for the same individuals, including contradictory records for gender, race, and birthdate. Notably, 1,779 subjects had inconsistent birthdates; in extreme cases, reported birthdates were 32 years apart, and in other instances, subjects’ reported ages decreased over time. Our analysis of the publicly available

MORPH-II [100], VisualBMI [14], FIW-BMI [26], and Illinois DOC [101] datasets further revealed that most recorded weights ended in digits of 0 or 5 (Fig S10). This pattern suggests that the weights were likely rounded from self-reported estimates. In contrast, the data collected from the Orang Asli, Tsimane, Ju/’hoansi, and Sama showed a uniform distribution across all terminal digits, closely matching the expected 10% frequency for each number. Establishing datasets from large-scale, market-integrated societies with reliable objective measurements is a prerequisite before we can accurately test cross-cultural generalizability or confidently deploy these tools for clinical or research purposes.

### Conclusion

Models including target populations in training data consistently outperformed those excluding it across all four populations, confirming that in-distribution training is preferred when possible. However, when target population data are unavailable, models trained on broader morphological diversity still outperform those trained on fewer populations. Global models achieved the lowest absolute errors because they captured continuous phenotypic variation across the groups; their cross-cultural predictions maintained accuracy and, for most populations, were nearly equivalent in absolute error to their focal counterparts.

Our findings have implications for the equitable training and deployment of ML tools in healthcare.

Models trained on datasets that lack cross-cultural representation risk encoding bias against the populations most reliant on remote and low-resource healthcare. Using more representative training data reduces this risk: our results provide a method for building models that generalize across diverse groups. Prioritizing training data collection across underrepresented global populations is therefore necessary for building medical ML tools that work reliably for everyone. When training data are sufficient, whether from broad cross-population coverage or from the focal population itself, these methods produce BMI estimates reliable enough for research use, such as generating retroactive longitudinal data or estimating BMI from photographs when anthropometric measurements are otherwise missing. However, a single model that performs reliably across all human populations remains out of reach without a greater diversity of training data.

## Acknowledgments

We thank the Orang Asli, Sama, Ju/’hoansi, and Tsimane participants who generously took part in this study. We also thank Timothy Webster, Shane Macfarlan, and Ryan Murdock (Advadnoun) who read and provided valuable feedback on earlier versions of this work.

## Supporting information

**S1 Fig.**
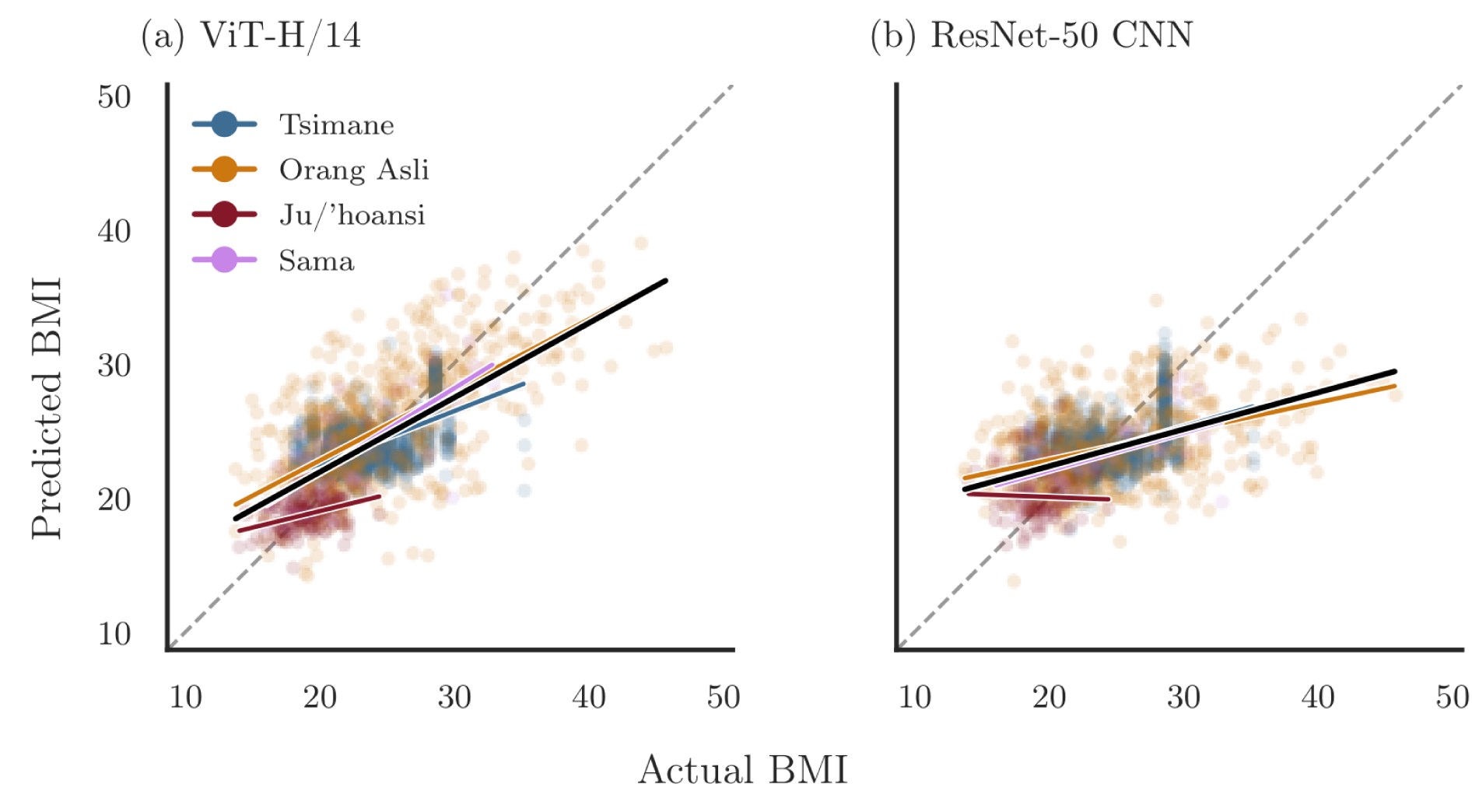
ViT versus CNN prediction accuracy. Actual versus predicted BMI values with our global model configuration generated by (a) the ViT-H/14 model and (b) the ResNet-50 CNN model.

**S2 Fig.**
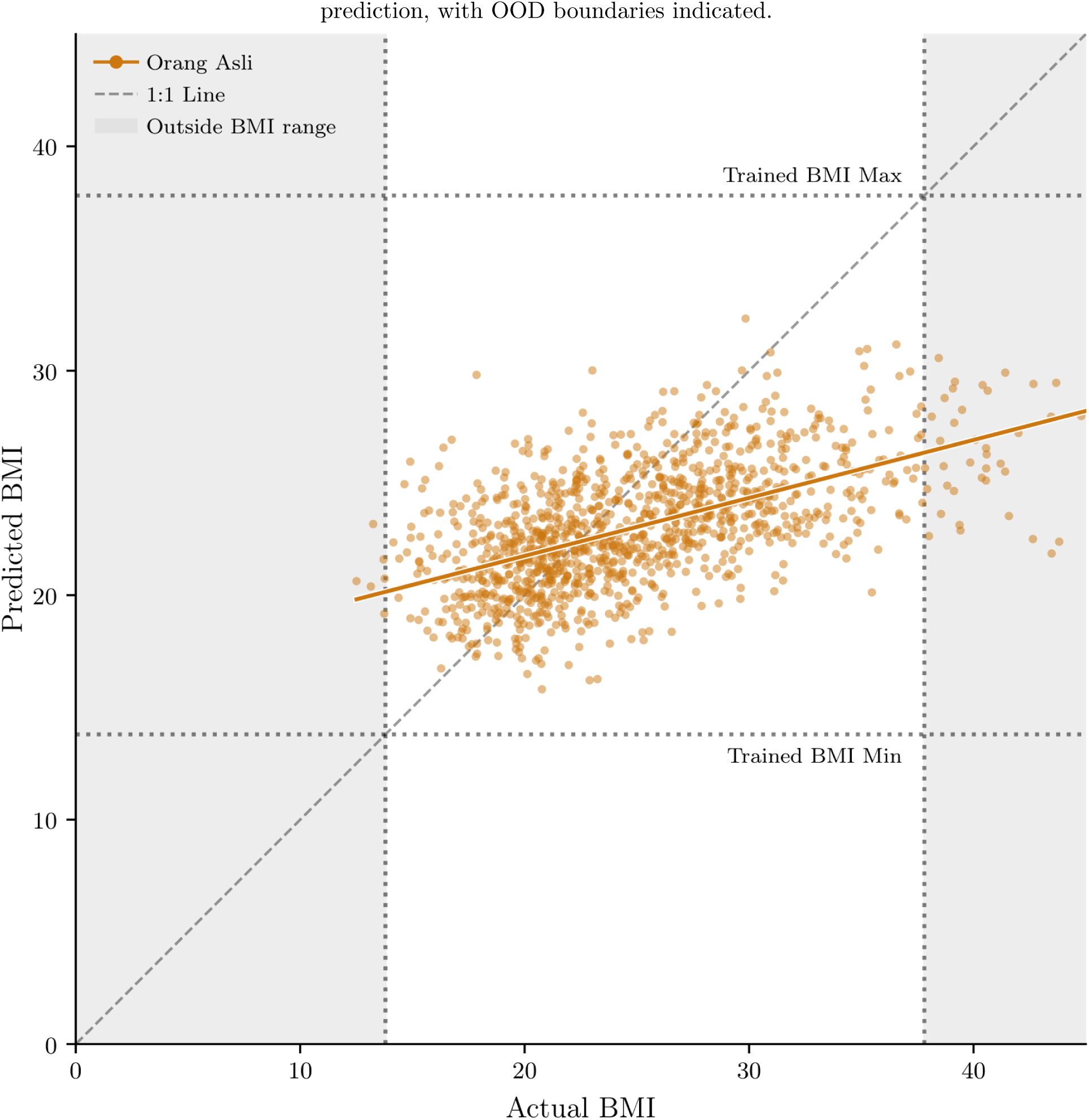
Out-of-distribution prediction boundaries. Scatter plot comparing actual and predicted BMI for the Orang Asli, including a linear regression trend line and a dashed 1:1 reference line for perfect prediction, with OOD boundaries indicated.

**S3 Fig.**
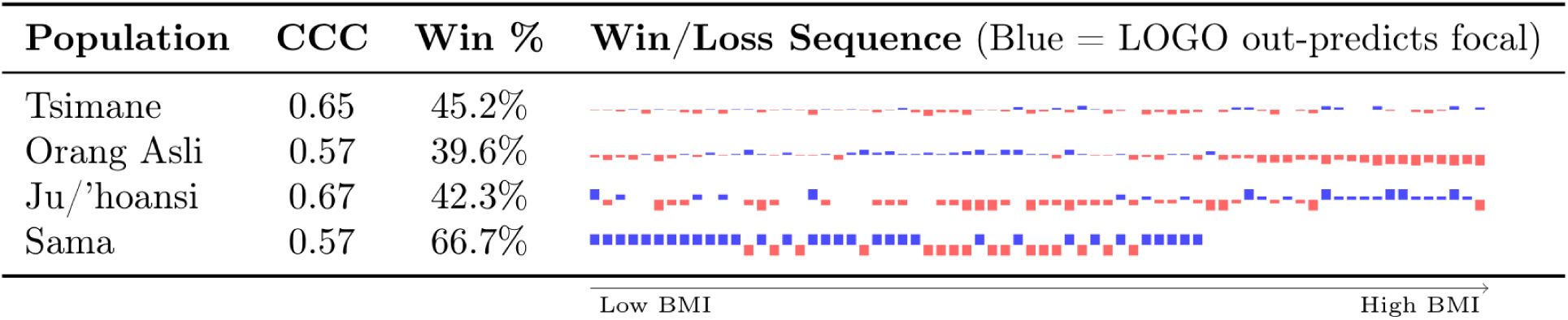
Evaluation of LOGO configuration performance against focal baselines. The CCC reflects the magnitude of structural agreement; the Win Rate indicates the overall percentage of samples where the LOGO model achieved a lower absolute error. The Win/Loss Sequence displays binned performance trends across each target group’s BMI spectrum (sorted lowest to highest): blue bars (pointing upward) denote BMI intervals where the LOGO model outperformed the focal model in the majority of cases, while red bars (pointing downward) indicate intervals of focal model superiority, with bar height reflecting the magnitude of the win rate within that interval.

**S4 Fig.**
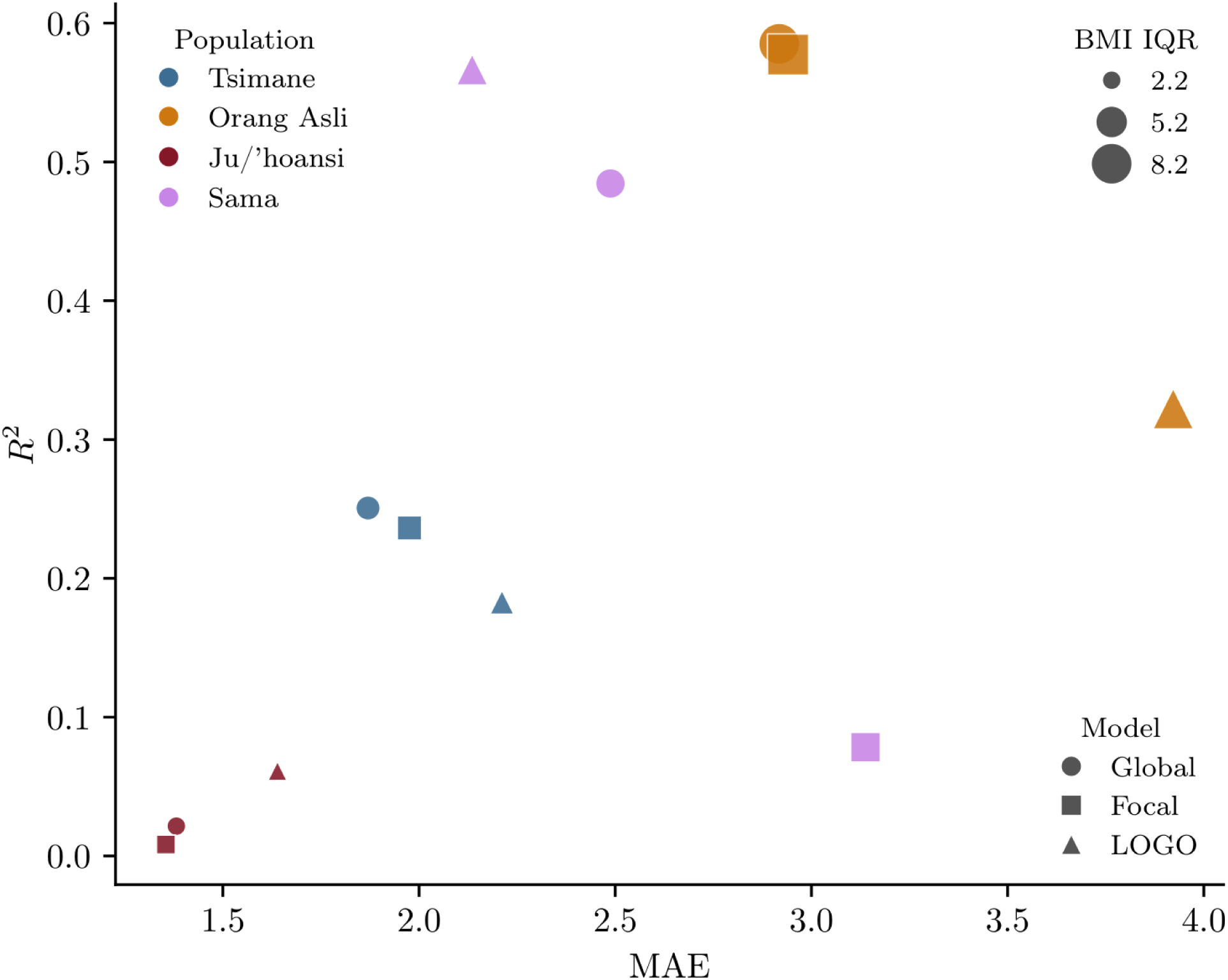
Relationship between MAE and *R*^2^ across population–model combinations.

**S5 Fig.**
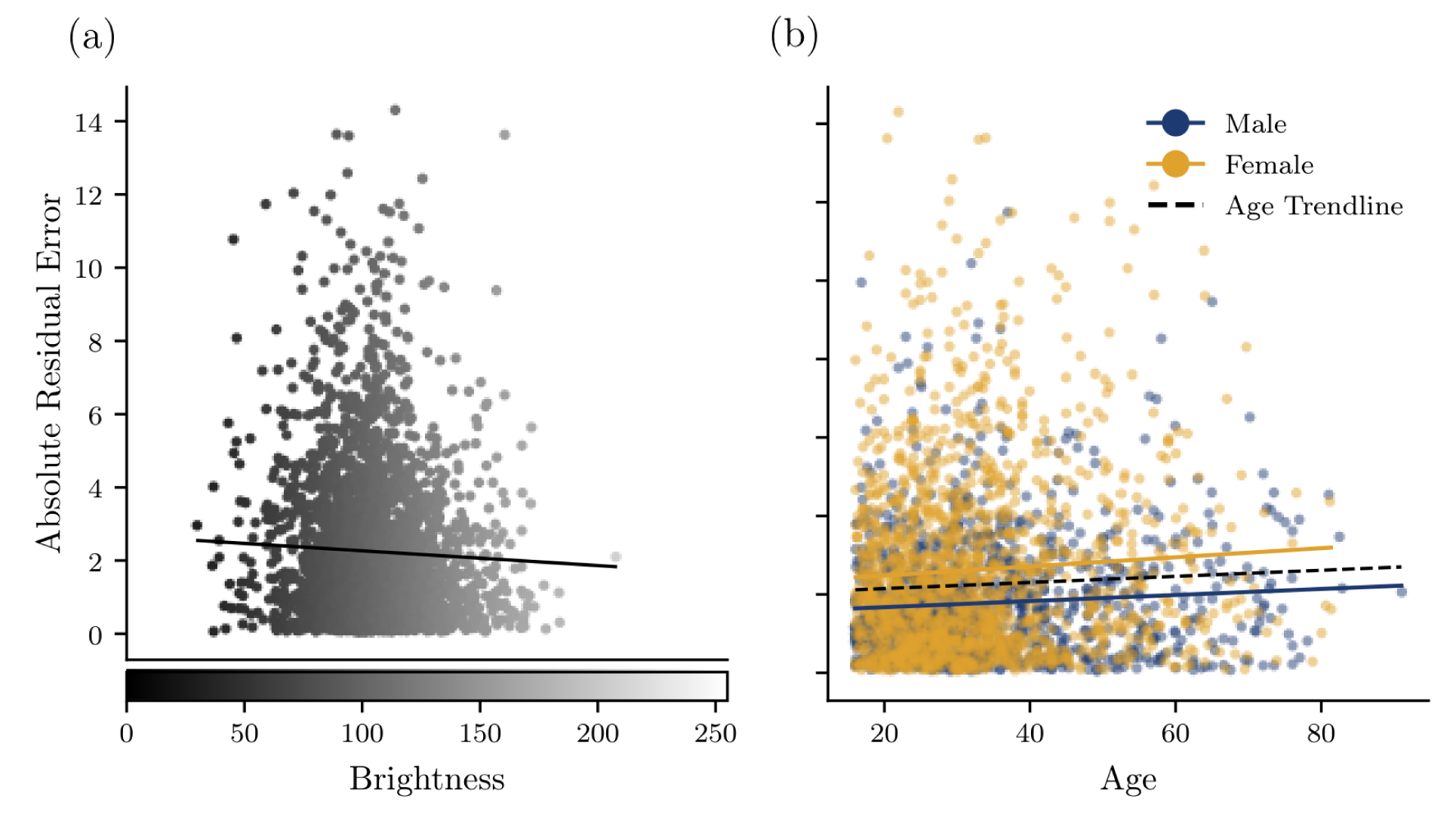
Absolute residual errors versus BMI deviation and demographic factors.

**S6 Fig.**
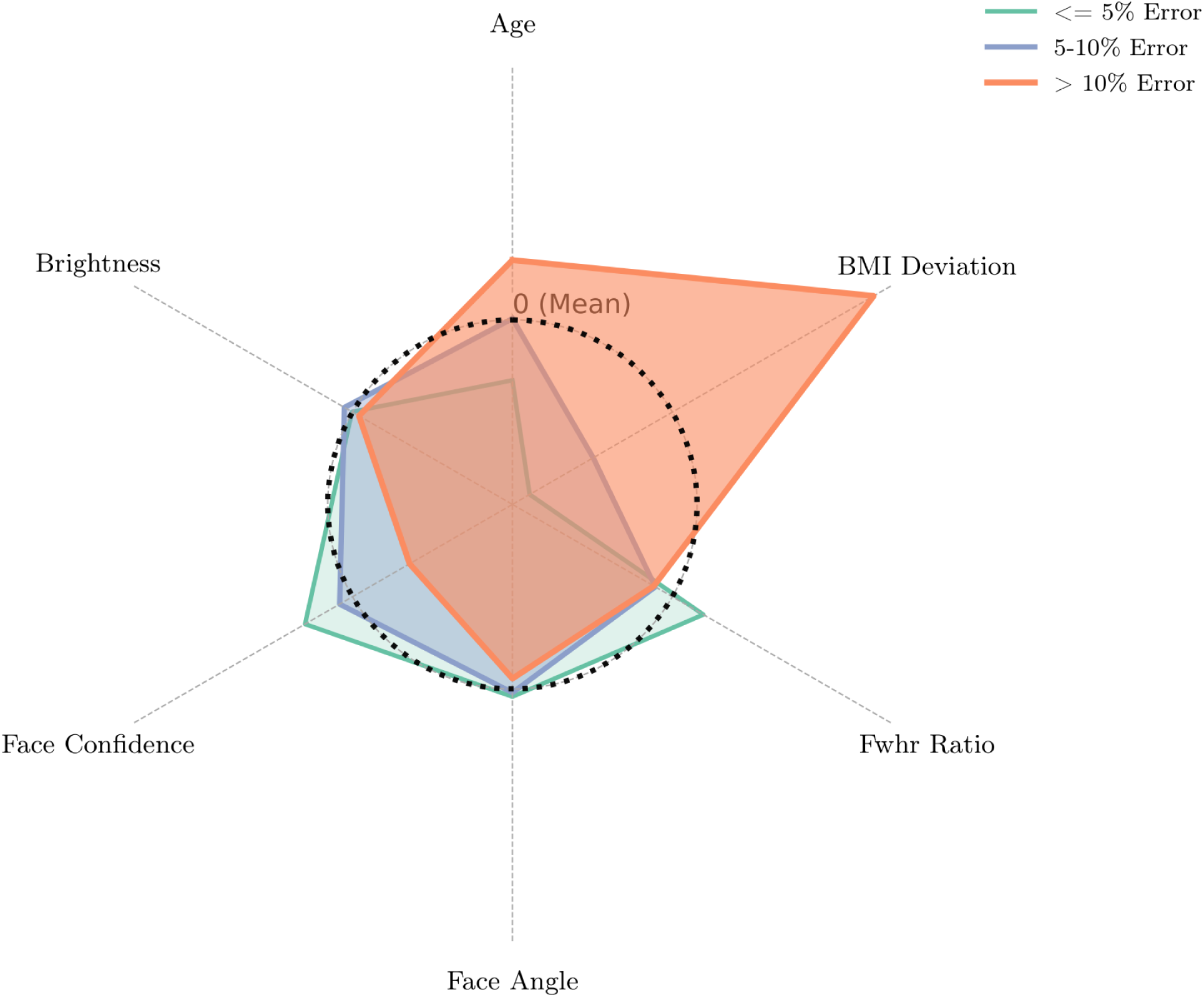
Demographic and biometric profiles of face-to-BMI predictions.

**S7 Fig.**
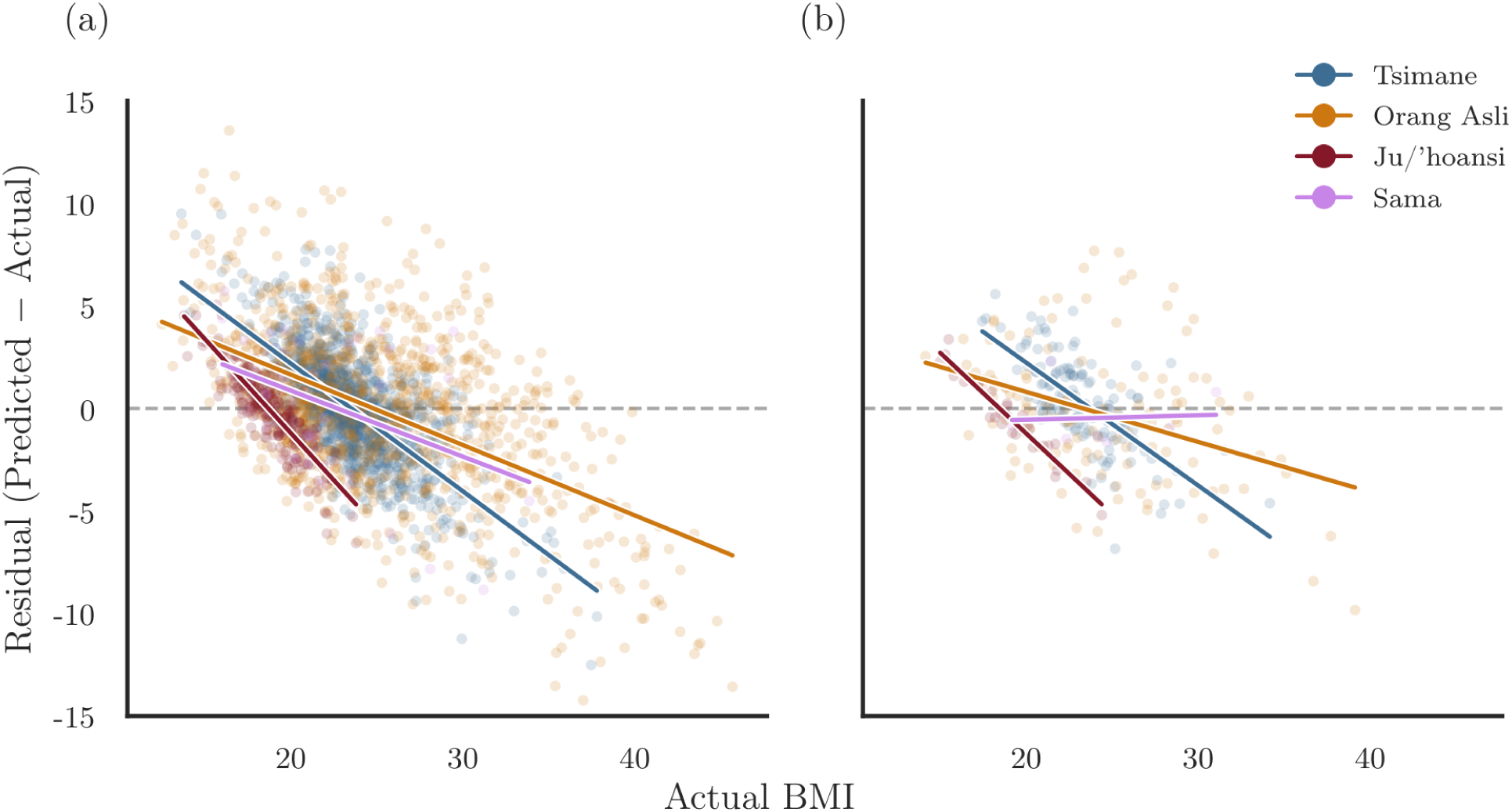
Global model prediction errors across the BMI spectrum for validation and test data.

**S8 Fig.**
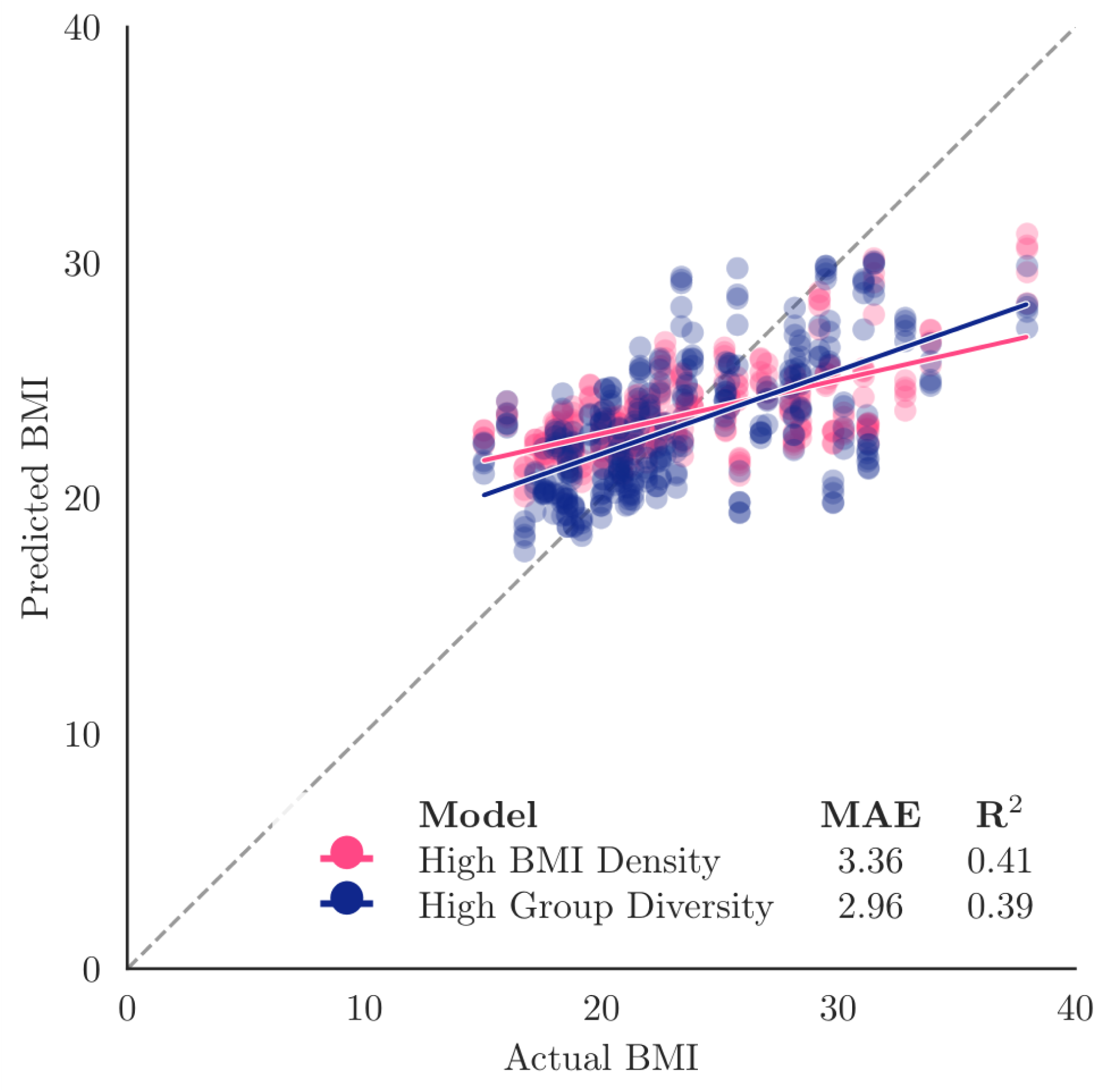
BMI training density versus morphological diversity. Comparison of two training strategies isolating the effects of BMI training density versus morphological variance, predicting the OOD Sama cohort.

**S9 Fig.**
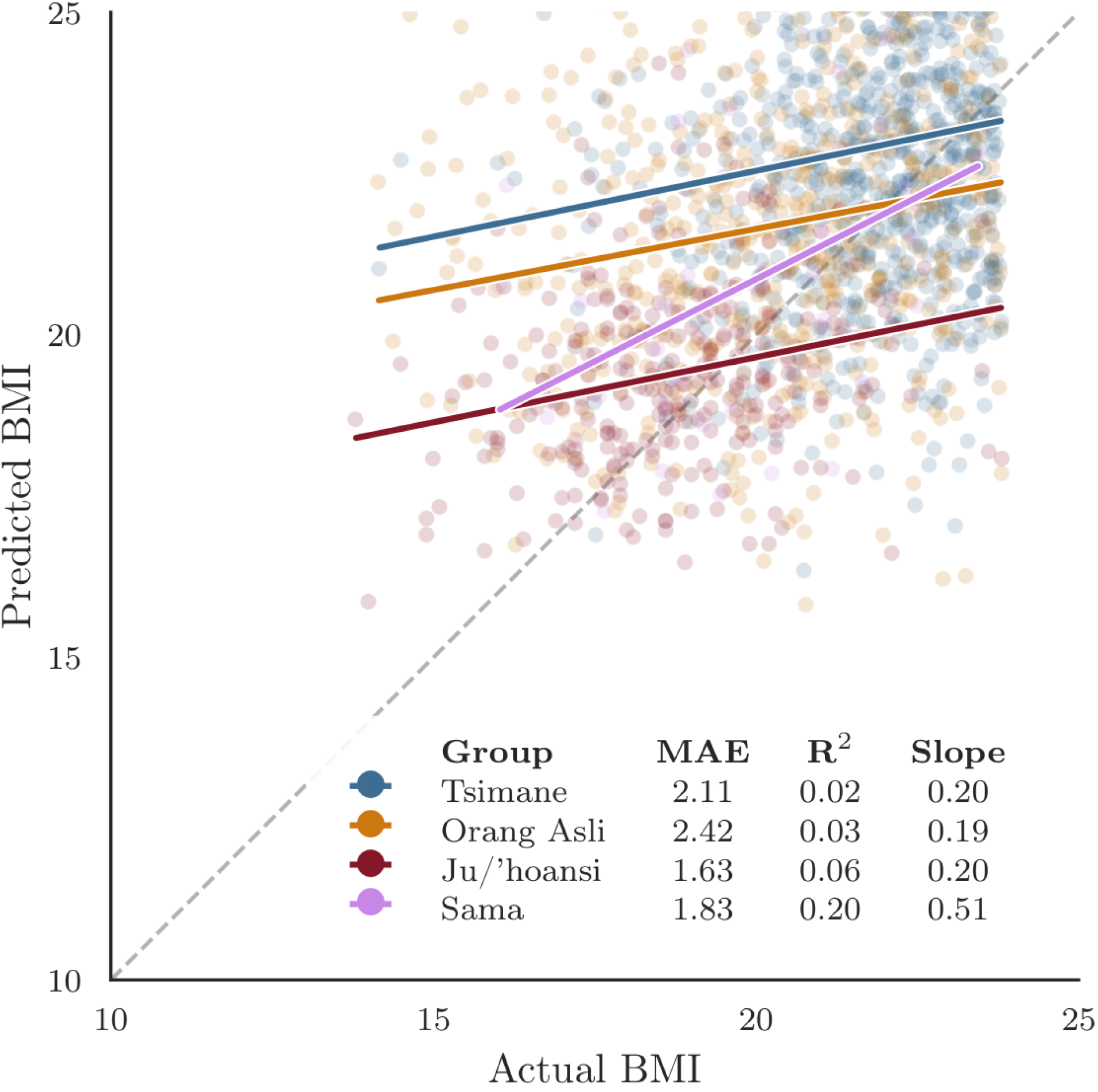
Effect of restricted BMI range on LOGO model regression slopes.

**S10 Fig.**
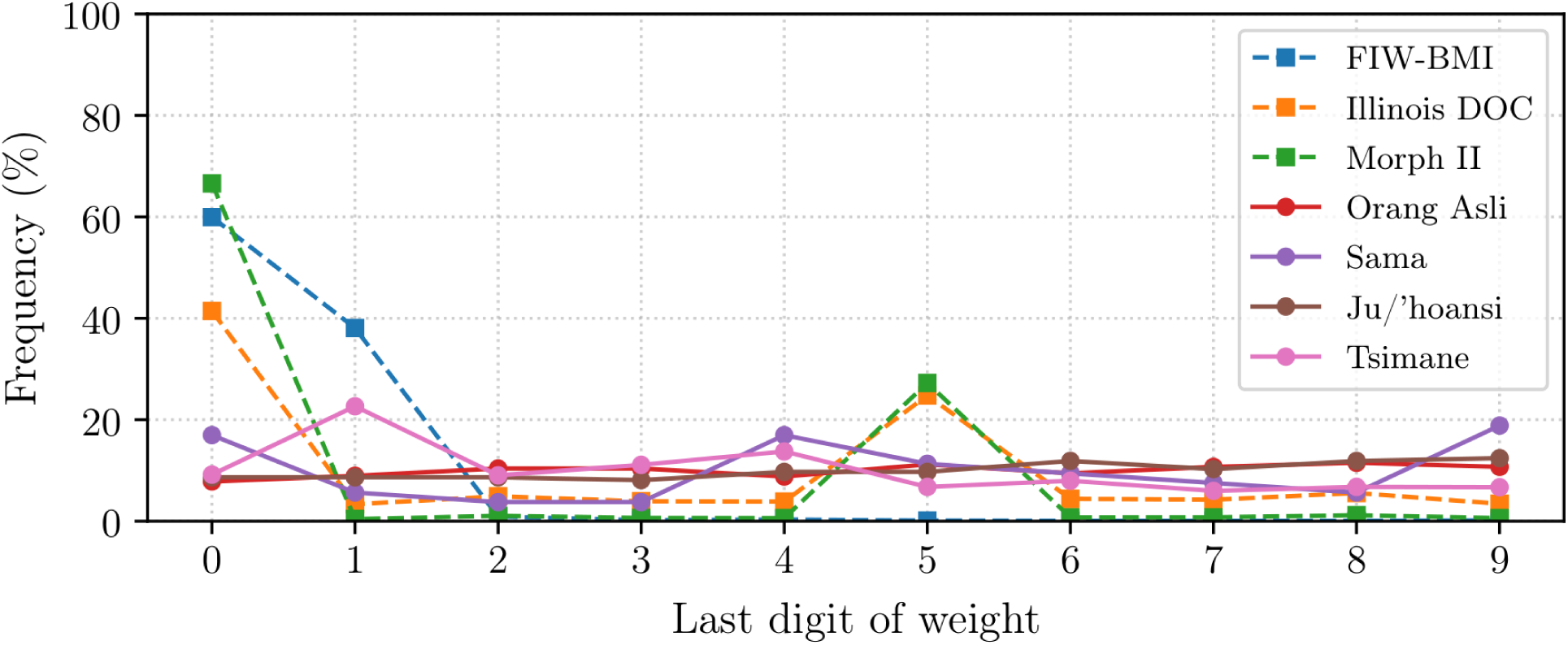
Distribution of the final digit of recorded weights across evaluated datasets.

**S1 Table.**
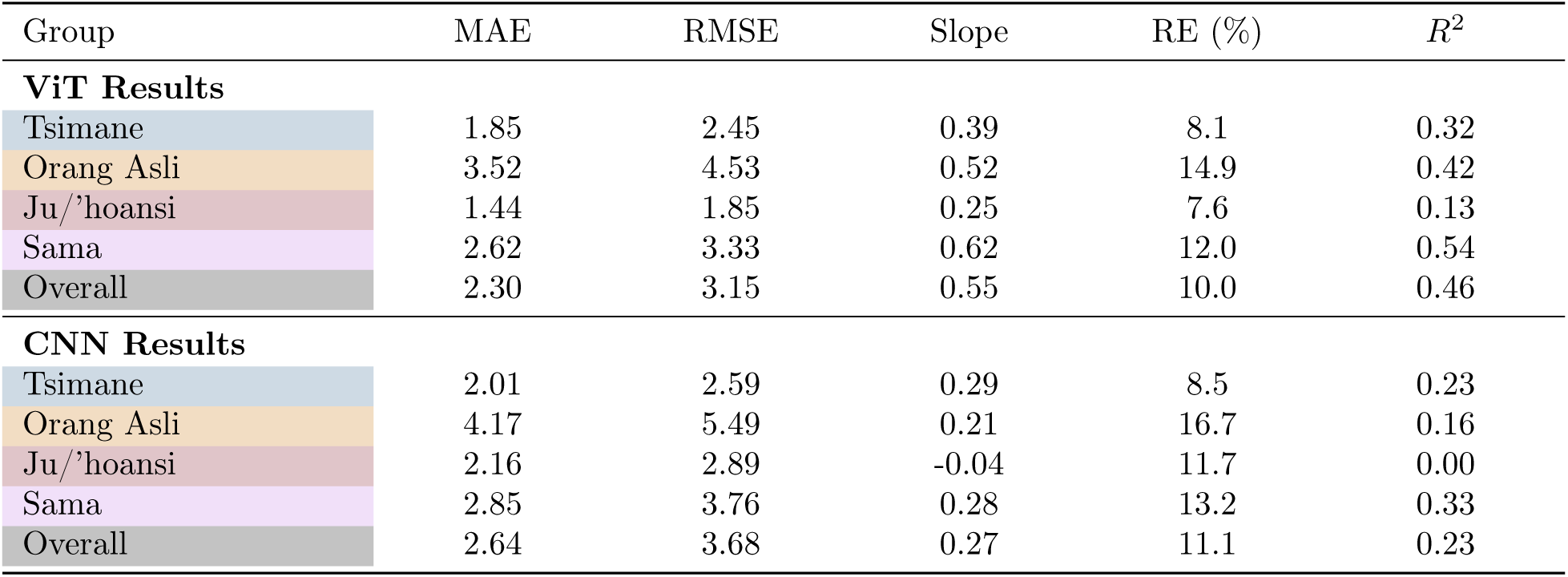
Comparison of absolute error metrics for the global model for the ViT and CNN predictions.

**S2 Table.** Evaluation of the global model across our four populations. For each group it reports the sample size, MAE, *R*^2^, WHO categorical accuracy, and the percentage of predictions within 5% and within 10% relative error.

| Group | $N$ | MAE | $R^2$ | WHO Acc. (%) | $\leq 5\%$ Err | $\leq 10\%$ Err |
| --- | --- | --- | --- | --- | --- | --- |
| Orang Asli | 1056 | 2.93 | 0.59 | 55.87 | 28.88 | 52.94 |
| Ju/'hoansi | 254 | 1.38 | 0.02 | 57.87 | 42.91 | 74.02 |
| Tsimane | 1301 | 1.87 | 0.25 | 73.79 | 42.20 | 70.33 |
| Sama | 48 | 2.49 | 0.48 | 58.33 | 20.83 | 50.00 |
| Overall | 2659 | 2.25 | 0.57 | 64.87 | 36.59 | 63.41 |

*Note:* WHO Acc. is the percentage of predictions falling in the exact correct WHO BMI category.

**S3 Table.**
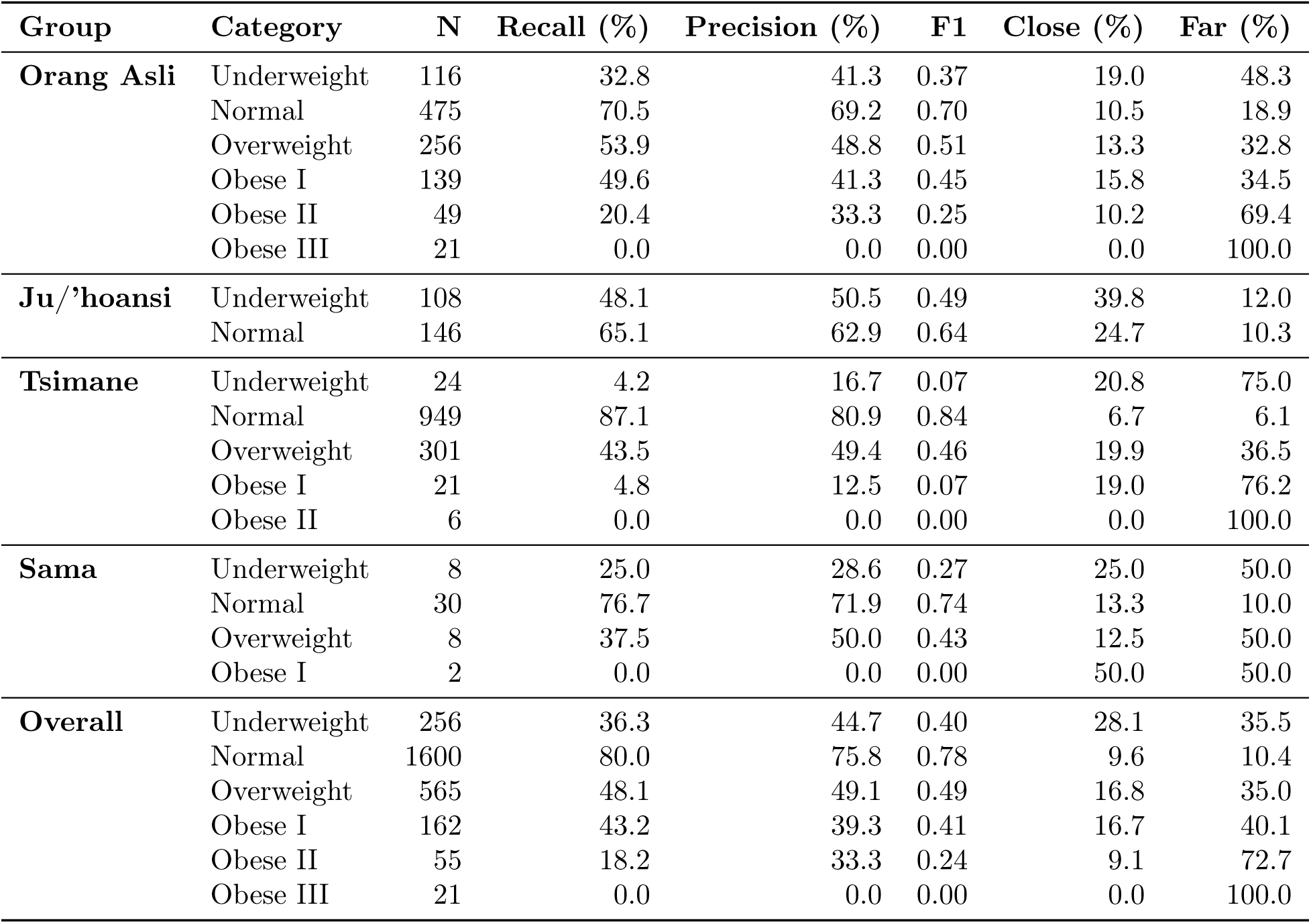
WHO category agreement in the global model. For each population and WHO BMI category, reports the number of individuals with that true category, recall, precision, F1 score, and the percentage of misclassifications that were close (one adjacent category) versus far (two or more categories away).

**S4 Table.** Percentage of BMI predictions within absolute relative error bands (*≤*5%, 5–10%, and >10%). Stratified by population group and model type (global, focal, LOGO), with an overall summary.

| Population | Global |  |  | Focal |  |  | LOGO |  |  |
| --- | --- | --- | --- | --- | --- | --- | --- | --- | --- |
| | $\leq 5\%$ | $5\text{--}10\%$ | $>10\%$ | $\leq 5\%$ | $5\text{--}10\%$ | $>10\%$ | $\leq 5\%$ | $5\text{--}10\%$ | $>10\%$ |
| Tsimane | 42.2 | 28.1 | 29.7 | 38.9 | 30.6 | 30.5 | 35.8 | 26.6 | 37.6 |
| Orang Asli | 28.9 | 24.1 | 47.1 | 27.9 | 24.2 | 47.8 | 19.8 | 19.4 | 60.8 |
| Ju/'hoansi | 42.9 | 31.1 | 26.0 | 40.6 | 36.2 | 23.2 | 36.3 | 29.2 | 34.5 |
| Sama | 20.8 | 29.2 | 50.0 | 27.1 | 8.3 | 64.6 | 39.3 | 21.4 | 39.3 |
| Overall | 36.6 | 26.8 | 36.6 | 34.5 | 28.2 | 37.3 | 29.6 | 23.9 | 46.6 |

